# Epigenetic motifs distinguishing endogenous from exogenous retroviral integrants

**DOI:** 10.1101/2025.04.30.651506

**Authors:** Sarah LaMere, Hanbei Xiong, Wei Wang, Brian LaMere, Niema Moshiri

**Affiliations:** Department of Medicine, University of California San Diego, La Jolla, California, USA; Department of Chemistry and Biochemistry, University of California San Diego, La Jolla, California, USA; Department of Cellular and Molecular Medicine, University of California San Diego, La Jolla, California, USA; Escondido, CA; Department of Computer Science and Engineering, University of California San Diego, La Jolla, California, USA

**Keywords:** DNA methylation, endogenous retrovirus, epigenetics

## Abstract

Retroviruses are subject to epigenetic regulation by the host genome after integrating, similar to vertebrate genes. However, their patterns of integration, and therefore their likely epigenetic regulation, differ between genera. Beta- and gammaretroviruses are two types of simple retroviruses that have a strong tendency to infect germ cells and endogenize. While ancient endogenous retroviruses are often easy to spot due to mutations rendering them non-functional, more recent integrants can maintain the capacity for full viral production, making it sometimes difficult to discern which integrants are exogenous and likely more clinically relevant. Because endogenous retroviruses generally spend a longer time integrated and subject to host epigenetic regulation as proviral DNA, we hypothesized we could show these integrants exhibit sequence differences from their exogenous counterparts, likely resulting from DNA methylation and histone modifications, and that endogenous retroviruses would generally show habituation to host promoters. Therefore, we have used statistical analyses of publicly available sequence data to demonstrate that endogenous retroviral variants exhibit decreased CpG dinucleotide and trinucleotide frequencies over time, and that they will show evidence for loss of motifs associated with ’active’ histone modifications. Close examination of these patterns provides further clues for distinguishing endogenous and exogenous retroviral variants, potentially aiding in the study of retroviruses in less well-characterized wildlife species.

**Importance:** Expression of vertebrate genes is regulated by chemical modifications made directly to the DNA or to the proteins associated with it, termed epigenetics. Because retroviruses integrate into DNA, they are subject to the same epigenetic modifications as regular genes.

Retroviruses will tend to endogenize, meaning they will become a permanent part of a species’ genome when their hidden DNA is passed down to progeny during reproduction. However, sometimes it is difficult to discern whether a retroviral sequence is endogenous (permanently fixed) or exogenous (an infectious entity). We hypothesized that changes to the retroviral sequences over time after endogenization would result from epigenetic modifications, and that these changes could help distinguish an endogenous retrovirus from an exogenous one. In this paper, we show that changes to the viral sequences associated with epigenetics indeed take place after endogenization.

## Introduction

Retroviruses are a unique family of viruses that are capable of integrating into the genomes of their hosts. Because these viruses carry their own distinctive polymerase capable of reverse transcribing the viral RNA genome into DNA, followed by integration into the host DNA, they are capable of permanently altering the genome in both the cell they infect and its progeny. When retroviruses infect germ cells, they can become incorporated into every cell of the resultant offspring, a process known as endogenization. Following endogenization, retroviruses will tend to acquire mutations rendering their promoter sequences and/or gene products deficient over the course of generations. Therefore, the degree of mutation of an endogenous retroviral sequence can give a window into how long it has been endogenized in a species.^1^ Over 40% of mammalian genomes are reported to be products of reverse transcription, including LINEs and SIINEs, with 8-10% derived from endogenous retroviruses.^2^

Retroviruses are divided into seven main genera based upon their genome organization and structures.^3^ These genera exhibit different propensities for the genomic regions into which they tend to integrate. For example, lentiviruses, which include human immunodeficiency virus (HIV) and feline immunodeficiency virus (FIV), have a tendency to integrate into transcriptionally active gene bodies. On the other hand, gammaretroviruses, which include murine leukemia virus (MuLV) and feline leukemia virus (FeLV), tend to target CpG-rich gene promoters.^4^ Betaretroviruses, such as mouse mammary tumor virus (MMTV) and Jaagsiekte sheep retrovirus (JSRV), have shown a weak propensity for transcriptional start sites (TSS),^5^ but generally seem to have a more random distribution of integration.^6^ Gammaretroviruses and betaretroviruses are much more prone than lentiviruses to infecting germ cells and endogenizing, as evidenced by their retroviral remnants being present in every major mammalian genome examined, including HERV-K (betaretrovirus) and HERV-W (gammaretrovirus) in humans. Interestingly, endogenous beta- and gammaretroviruses can frequently have exogenous counterparts that are fully infectious and replication-competent.

Their sequences are often only slightly divergent from each other, usually in the long terminal repeat (LTR) and/or envelope (env), impacting their expression and tropism, respectively.^7^

Endogenous and exogenous retroviral variants are well-characterized for some species, such as MuLV and FeLV. However, the distinction between recent integrants with transcriptionally intact genomes in more poorly characterized species can be difficult. A case in point is the koala retrovirus, KoRV, which is a gammaretrovirus in the process of endogenizing. The virus has been associated with leukemias and lymphomas, as well as with secondary infections such as chlamydia.^8^ Thus far, ten different subtypes have been described, varying with geography and exhibiting different associations with disease. KoRV-A appears to be the endogenous variant that has arisen from this virus, while the other subtypes exhibit more overt associations with disease and are believed to be exogenous.^9^

To make matters more confusing, endogenous and exogenous retroviruses will recombine. The endogenous variant of FeLV can recombine with the exogenous variant FeLV-A to yield a new virus with different tropism, and recombination events can even occur with other types of less related endogenous retroviruses, giving rise to diversity and evolution within the genera.^10^ Most of these recombination events result from viruses within a single or closely related species, but there are examples of recombination between gammaretroviruses from more divergent species. For example, RD114, another endogenous gammaretrovirus found in domestic cats, appears to be the product of a recombination event between an endogenous feline retrovirus (*Felis catus* endogenous retrovirus, or FcEV) and baboon endogenous retrovirus (BaEV) and is believed to have resulted from a cross-species viral transfer from an old-world monkey to a domestic cat.^11^

Multiple host epigenetic mechanisms used to suppress endogenous retroviral infections have been described, including DNA methylation, histone modifications, and non-coding RNAs.^12^ Exogenous retroviruses can also be subject to these same mechanisms as a molecular arm of the host antiviral defense, making them an active area of study in HIV.^13^ However, endogenous retroviruses that have co-evolved with their host species are frequently more highly transcriptionally suppressed than their exogenous counterparts,^7^ prompting the hypothesis that sequence-specific evidence of epigenetic suppression will generally be more apparent in endogenous variants.

Previously, HIV was reported to mirror the human genome with its low CpG content, exhibiting evidence that this was primarily a product of deamination following CpG methylation rather than other possible mechanisms, such as CpG depletion by APOBEC3 or due to toll-like receptors.^14^ Because of their propensity to integrate into promoter regions, we hypothesized that gammaretroviruses would mirror the CpG content of the promoters in their respective species, and originally, we also hypothesized that CpG content of betaretroviral genomes might mirror that of promoters due to a TSS integration bias. We also hypothesized that endogenous variants would exhibit decreased CpG content relative to exogenous variants, suggesting more CpG methylation and deamination following endogenization, as well as altered CpG trinucleotide motifs consistent with the methylation patterns in their host species. Finally, we hypothesized that motifs consistent with active histone modifications would be more apparent in exogenous proximal proviral sequences, while those more consistent with repressive modifications (e.g. H3K9me3) would be more evident in known endogenous sequences. To test these hypotheses, we performed statistical analyses on CpG frequencies and CpG trinucleotide motif frequencies from various retroviruses and compared endogenous to exogenous variants, as well as all those from promoter sequences of the species in which they are present. We also performed bisulfite mapping to examine methylation patterns in endogenous retroviruses versus promoters. Finally, we examined differences in motifs associated with histone modifications using the Simple Enrichment Analysis tool from the MEME suite.^15^ Because betaretroviruses have different integration propensities than gammaretroviruses, we focused on characteristics for the few viruses we were able to examine in each genus, as their integration behavior potentially impacts their epigenetic regulation.

## Results

### Recently endogenized beta- and gammaretroviruses reflect CpG content of eutherian host promoters, with variability attributable to DNA methylation

To evaluate how betaretroviruses and gammaretroviruses adapt to the genomes of their respective species, we examined CpG content of full-length endogenous and exogenous retroviruses from cats, mice, pigs, sheep, and koalas from **Tables 1 and 2** (see **Data Availability Statement** for accession numbers) using a Python-based Kmer tool. The Kmer tool calculates D-ratios, which are the observed/expected (or O/E ratios) based on a first order Markov model of conditional probabilities (see **Methods**). We assessed the CpG content of endogenous retroviruses relative to multiple genomic regions (500 bp upstream of the TSS, 1kb upstream of the TSS, 2kb upstream of the TSS, whole gene bodies, 1kb downstream of all genes, and entire chromosomes) in all five species **(Supplementary Figures S1 and S2)**.

**Table 1:** Endogenous with counterpart exogenous retroviruses used for analysis.

| Gammaretroviruses |  |  |  |  |  |  |
| --- | --- | --- | --- | --- | --- | --- |
|  | Exogenous<br>vs<br>endogenous<br>% identity | Original<br>Integration<br>time | Exogenous CpG<br>O/E | Endogenous<br>CpG O/E | Exog. N | Endog. N |
| FeLV | 86% | <2 MYA<sup>16</sup> | 0.53 | 0.49 | 8 | 8 |
| MuLV | 90-100% | 1 MYA<sup>17</sup> | 0.53 | 0.52 | 16 | 49 |
| KoRV | 90% | 22,000-49,000<br>YA<sup>18</sup> | 0.51 | 0.5 | 8 | 16 |
| Betaretroviruses |  |  |  |  |  |  |
| MMTV | 95-99% | 20 MYA<sup>19</sup> | 0.45 | 0.46 | 6 | 5 |
| JSRV | 90-98% | 5-7 MYA<sup>20</sup> | 0.58 | 0.57 | 20 | 19 |

**Table 2:**
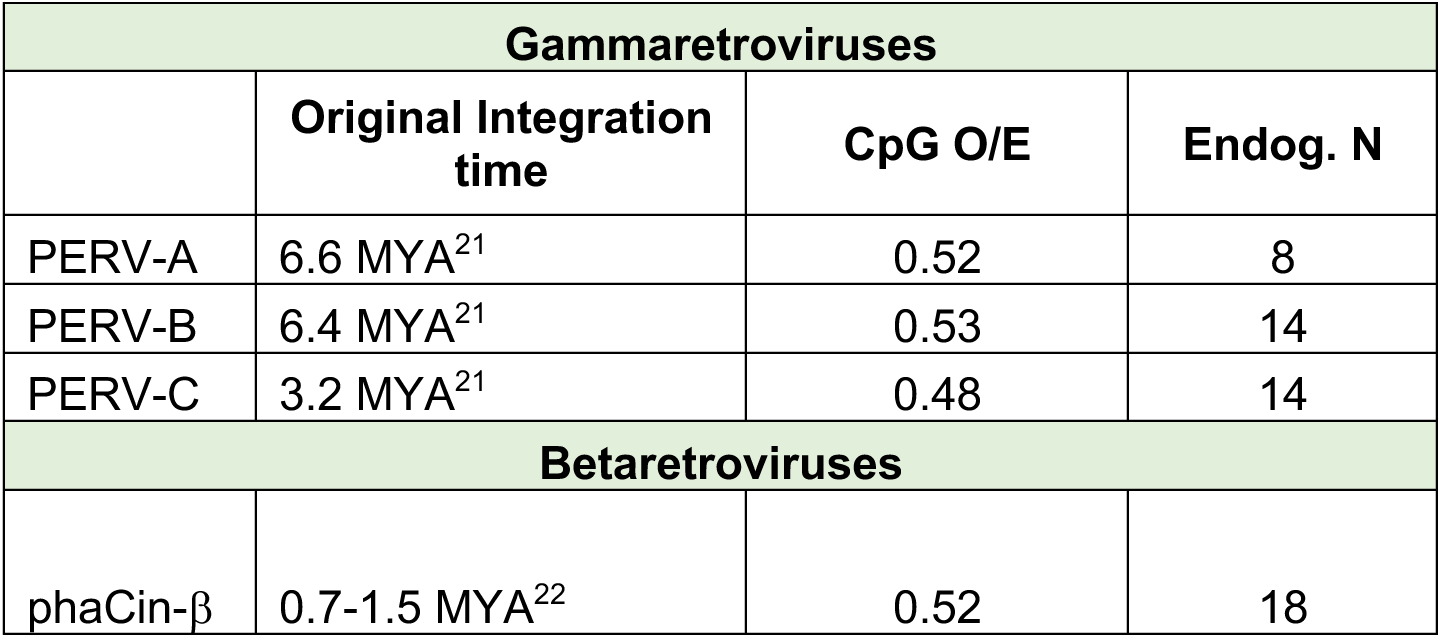
Further endogenous retroviruses used for analysis.

In most cases, ERVs reflected the CpG content of regions 500 bp upstream of the TSS, especially for the gammaretroviruses we evaluated **(Supplementary Figure S1a-d).** The betaretroviruses we evaluated also trended this way, although enJSRV in particular had a higher CpG O/E ratio relative to the other endogenous retroviruses we examined **(Supplementary Figure S2a-c)**. However, the five different mammalian species exhibited significantly different CpG O/E ratios from each other in promoter regions (defined in **Supplementary S2d** as 1 kb upstream of the TSS), suggesting that while endogenous beta- and gammaretroviruses averaged CpG O/E ratios around 0.5, these could deviate significantly from promoter CpG content, depending on the species they infect. Cats in particular had overall higher CpG content across all regions examined **(Supplementary Figure S1a**, **Supplementary Figure S2d)**, while sheep, koalas, and mice exhibited lower relative CpG content closer to 0.4 within 1kb of the TSS **(Supplementary Figure S2d)**.

Based on our analysis, we compared CpG content of endogenous and exogenous beta- and gammaretroviruses in each species to that of the region within 500 bp upstream of the TSS **(Figure 1a-e)**. Mice and sheep both displayed significant differences in CpG content of their promoters from their respective retroviruses, reflecting their overall lower promoter CpG content. However, three out of the five species examined had no significant difference between their retroviral CpG content and their promoters. To rule out the potential impact of ancient retroviral sequences inhabiting promoters, we used also LTRHarvest^23^ to remove all LTR and retrotransposon-like sequences and re-analyzed CpG content using the Kmer tool, which resulted in a minimal change to the CpG content of the analyzed promoter sequences (data not shown).

**Figure 1:**
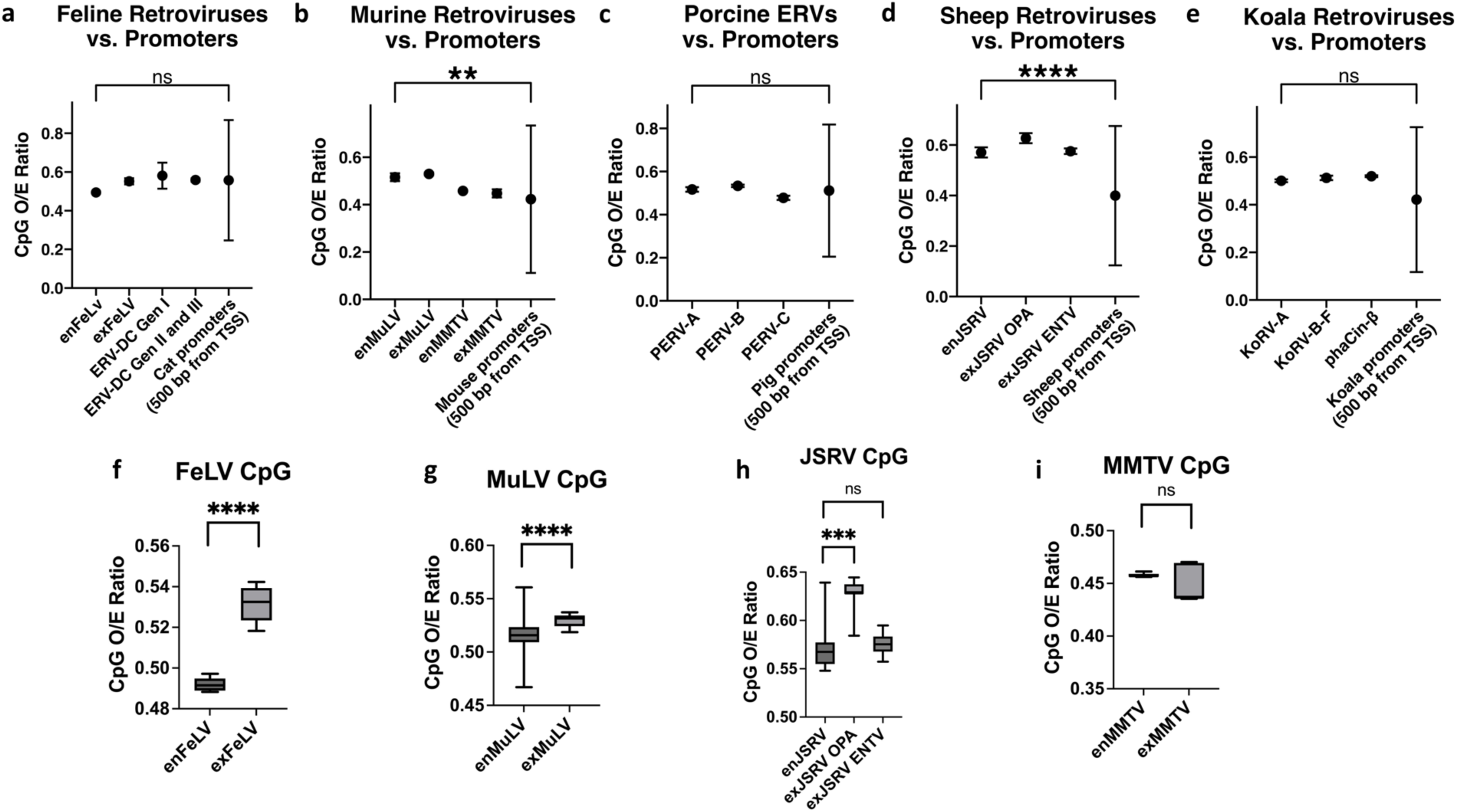
CpG content of beta- and gammaretroviruses commonly reflects CpG content of promoter sequences, and CpG content is usually lower with endogenization. **a-e)** Endogenous and exogenous retroviral median CpG D-ratios are compared to those of promoters (500 bp upstream of the TSS) for **a)** domestic cats **b)** mice **c)** pigs **d)** sheep and **e)** koalas. **f-h)** Endogenized retroviral variants exhibit lower CpG content than exogenous counterparts for **f)** Feline Leukemia Virus (FeLV), **g)** Murine Leukemia Virus (MuLV), and **h)** Jaagsiekte sheep retrovirus (JSRV). **i)** Mouse Mammary Tumor Virus (MMTV) shows no significant change to median CpG content. P-values are reported from Kruskal-Wallis tests for multiple comparisons **(a-e)** and Mann-Whitney comparisons **(f-i)**. Error bars represent interquartile ranges. n.s. p > 0.05, *** p < 0.001, **** p < 0.0001

### Endogenization impacts CpG content of beta- and gammaretroviruses due to CpG methylation

When comparing CpG content of endogenous vs. exogenous retroviruses, the endogenous gammaretroviruses FeLV and MuLV exhibited a statistically significant decrease in CpG content compared to their exogenous counterparts (**Figure 1f-g**). However, the betaretroviruses analyzed (MMTV and JSRV) did not have as robust a difference with the sequences available **(Figure 1h-i)**. In fact, no significant difference was noted for endogenous vs. exogenous MMTV, but the distribution of CpG content for MMTV is highly skewed due to very few full-length replicates being publicly available (i.e. 5 for enMMTV and 6 for exMMTV), making concrete conclusions difficult to draw. However, JSRV had many more replicates available, with 19 replicates for endogenous JSRV and 20 replicates for exogenous JSRV. JSRV has two exogenous variants, one which causes enzootic nasal tumor virus (ENTV), and another which causes ovine pulmonary adenocarcinoma (OPA).

Only the ENTV version of JSRV differed significantly in CpG content compared to the endogenous variant, despite our own phylogenetic analysis of full-length sequences suggesting it to be more closely related to the endogenous JSRV variant than the exogenous OPA variant is (**Supplementary Figure S3**).

To examine the role of DNA methylation in the CpG content of all retroviruses examined, we calculated the correlation between CpG, TpG, and CpA O/E ratio to assess for deamination of methylated cytosines. Deamination of methylated cytosines occurs over time, changing the CpG landscape to TpG and CpA on the complementary strand, while non-methylated cytosines are not subject to these same forces.^24^ Strikingly, both the endogenous and exogenous JSRV showed a strong significant symmetrical negative correlation **(Figure 2a-c)**. The correlation for exogenous JSRV was surprisingly more profound than that found in the endogenous variant. Of the endogenous retroviruses examined that had a closely related exogenous variant, JSRV has been one of the longest circulating, prompting speculation that perhaps long-standing integrants have relaxed epigenetic control. However, endogenous JSRV showed heavy methylation across the provirus, with a preference for CGC and CGG motifs **(Figure 2d-e)**. Consistent with the predilection for CGC methylation, CGC motifs had the strongest correlation with TGC/CAC **(Figure 2f)**, while endogenous JSRV showed a significant decrease in CGC O/E ratios compared to both exogenous JSRV subtypes **(Figure 2g)**. MMTV had few full-length replicates, three of which had identical sequences, calling into question the statistical power of the analysis for this virus. However, Bismark bisulfite mapping also demonstrated heavy CpG methylation across the body of the endogenous MMTV provirus, with CGC trinucleotides exhibiting a small but significantly higher percent methylation **(Figure 2h-i)**. CGC O/E ratios did not exhibit significant differences between endogenous and exogenous MMTV (data not shown), but few replicates were available for this analysis, making it difficult to achieve significance.

**Figure 2:**
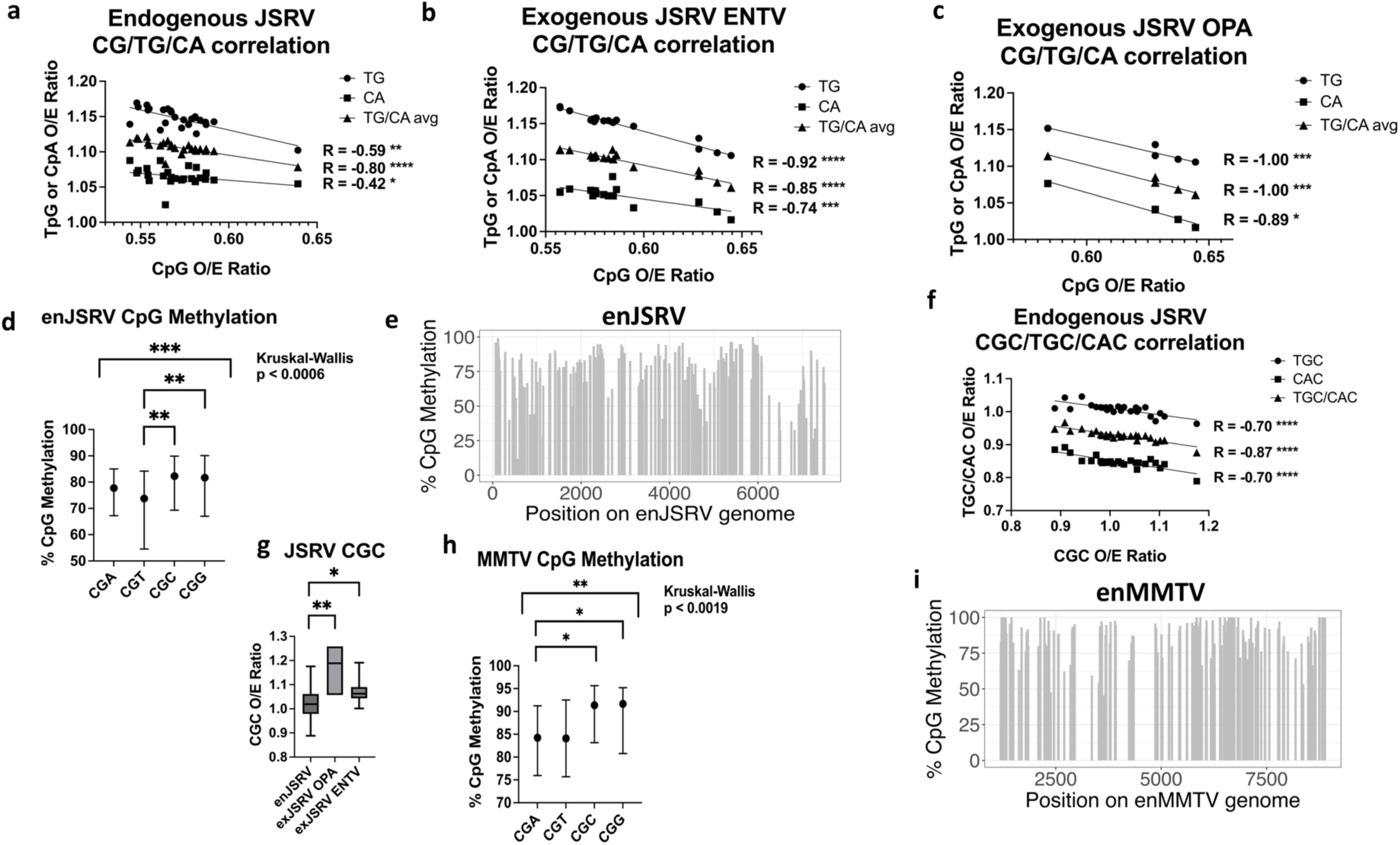
Endogenous betaretroviruses JSRV and MMTV are highly methylated and exhibit CpG decline. **a-c)** JSRV shows a strong negative correlation between CpG content and TpG/CpA content in **a)** endogenous **b)** exogenous Enzootic Nasal Tumor Virus (ENTV) and **c)** exogenous Ovine Pulmonary Adenocarcinoma (OPA) variants. **d)** CpG methylation across endogenous JSRV shows uneven CpG trinucleotide distribution, with a preference for CGC and CGG trinucleotides. **e)** Bismark mapping shows heavy CpG methylation across the entire JSRV provirus. **f)** Endogenous JSRV shows a strong negative correlation between CGC content and TGC/CAC. **g)** CGC content is significantly decreased in endogenous JSRV compared to both exogenous JSRV strains. **h)** Trinucleotide analysis of MMTV methylation shows a significant preference for CGC motifs. **i)** Bismark mapping shows heavy CpG methylation across the body of the MMTV provirus. R and p-values for correlation analyses are from Spearman correlations **(a-c,f)**. P-values are reported for the Kruskal-Wallis test for multiple comparisons with Dunn’s test **(d,g,h)**, while error bars in **d)**, **g)**, and **h)** represent interquartile ranges. *p < 0.05, **p < 0.01, ***p < 0.001, ****p < 0.0001

Correlations between CpG and TpG/CpA in the two gammaretroviruses we examined were less pronounced; FeLV, which only included 8 replicates of endogenous and exogenous full-length sequences each that were publicly available, showed no significant negative correlation, and differences in TpG and CpA O/E ratios were not significant between endogenous and exogenous variants. However, O/E ratios of the trinucleotide motif CGC showed a significant decrease in enFeLV compared to exFeLV, while TGC and CAC were significantly increased **(Figure 3a)**, suggesting this trinucleotide motif in particular is prone to DNA methylation in the context of enFeLV. Bisulfite mapping to cat promoters (defined as 1kb upstream of all transcriptional start sites) showed significantly lower CGC and CGG methylation **(Figure 3b)**, suggesting this pattern is inconsistent with that seen in cat promoters. Unfortunately, mapping the bisulfite converted cat genome to enFeLV failed to produce robust results, preventing definitive conclusions about the methylation patterns in enFeLV and their relationship to that of promoters in the domestic cat.

**Figure 3:**
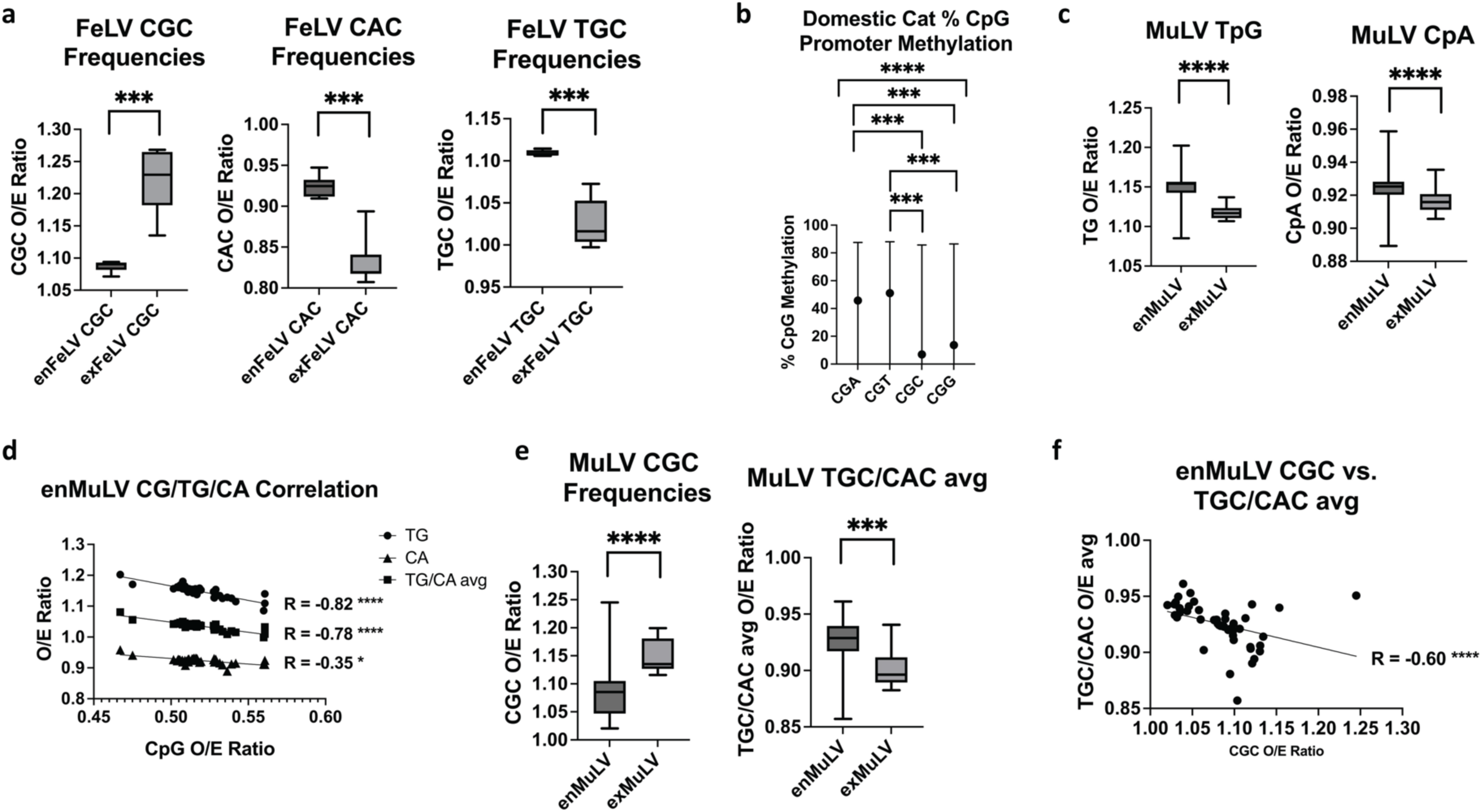
Endogenization of gammaretroviruses FeLV and MuLV impacts retroviral CpG content: **a)** FeLV endogenization leads to a significant decrease in CGC content, with corresponding increases in CAC and TGC content. **b)** Domestic cat promoters exhibit uneven median CpG methylation across trinucleotides, with CGC being the least methylated. **c)** Endogenous MuLV exhibits significantly increased TpG and CpA content compared to exogenous variants. **d)** enMuLV shows a strong negative correlation between CpG, TpG, and CpA, suggesting methylated cytosine deamination of enMuLV. **e)** EnMuLV CGC frequencies are significantly decreased compared to exMuLV, with corresponding increases in TGC/CAC frequencies. **f)** enMuLV shows a strong negative correlation between CGC and TGC/CAC, suggesting methylated cytosine deamination of enMuLV at CGC trinucleotides. P-values are reported for Mann-Whitney tests **(a, c, e)**, Kruskal-Wallis test for multiple comparisons **(b)**, and Spearman correlation **(d, f)**. Error bars for **b)** represent interquartile ranges. *p < 0.05, **p < 0.01, ***p < 0.001, ****p < 0.0001

More replicate sequences of endogenous and exogenous MuLV were available for analysis from ^25^, allowing a comprehensive examination of this gammaretrovirus. We analyzed promoters from three endogenous variants: polytropic (PMV), modified polytropic (MPMV), and xenotropic (XMV) MuLV. In addition to a significant decrease in CpG O/E ratios between enMuLV and exMuLV **(Figure 1g)**, enMuLV also exhibited a significant increase in TpG and CpA O/E compared to exMuLV, suggesting a likely role by DNA methylation **(Figure 3c)**. A significant correlation was also observed between CpG O/E and TpG/CpA O/E ratios in enMuLV, providing further support for this conclusion **(Figure 3d)**. Bisulfite analysis of mouse promoters showed no significant difference in frequency of trinucleotide motifs methylated in mouse promoters, and enMuLV also showed no difference **(Supplementary Figure S4)**.

CGG trinucleotides showed no significant difference in frequency between enMuLV and exMuLV, and while CCG trinucleotides did show a difference, CCG frequency negatively correlated only with CCA and not CTG **(Supplementary Figure S5a-e)**, suggesting APOBEC3G was a more likely cause of the asymmetric deamination.^26^ Analysis of all enMuLV sequences using Hypermut 3^27^ demonstrated significant hypermutation due to APOBEC3G for all but three of the 49 sequences examined **(Supplementary Table S1)**, providing further support for this. Of note, the paper from which these enMuLV sequences were retrieved discusses the role of APOBEC3 in genetic diversity among enMuLVs.^25^ Interestingly, as was true for FeLV, a significant reduction was found in CGC trinucleotide motifs in the endogenous MuLVs compared to exMuLVs, while a significant increase in corresponding TGC and CAC frequency was also observed **(Figure 3e)**. A modest but significant negative correlation between CGC with TGC and CAC was also found in enMuLV **(Figure 3f)**, supporting the CGC motif as a more frequent past target for DNA methyltransferases in enMuLV. Unlike the methylation patterns in the other endogenous retroviruses, MuLV did not exhibit a significant preference for CGC methylation, nor did mouse promoters **(Supplementary Figure S4)**, calling into question whether CpG methylation is responsible in this case. However, these CpG frequency patterns might reflect initial methylation following integration rather than following endogenization. In all cases comparing endogenous and exogenous variants, there was evidence of a predilection for CGC methylation following endogenization, either via available methylation data, CGC O/E data, or both.

### CpG content of endogenous retroviruses is variably impacted by DNA methylation and corresponds with transcriptional activity

Pigs have three highly related endogenous gammaretroviruses known as Porcine Endogenous Retrovirus A, B, and C. The viral sequences are primarily different in their envelope regions, as well as some difference in the LTRs that appear to confer varying levels of transcriptional activity. PERV-A is the oldest, having endogenized approximately 6.6 million years ago. PERV-B is close in age, having endogenized approximately 6.4 million years ago.

Finally, PERV-C is the most divergent of the three and is thought to have endogenized 3.2 million years ago.^28^ Of the three, PERV-B is the most transcriptionally active, while PERV-C is the least, and the CpG contents of the three proviruses reflect this, with PERV-B exhibiting the highest CpG O/E ratio, and PERV-C exhibiting the lowest **(Figure 4a)**. Of the three proviruses, only PERV-C demonstrates a correlation between CpG and TpG/CpA O/E ratios **(Figure 4b)**, suggesting this provirus is the most heavily impacted by DNA methylation. To attempt to confirm this finding, we mapped a bisulfite converted pig genome to full-length PERV-A, PERV-B, and PERV-C. The PERV-C envelope had no reads mapping, suggesting this variant was not present in the publicly available bisulfite converted pig genome we accessed. However, methylation was heavy across both PERV-A and PERV-B **(Figure 4c)**. Because sequences are identical for the two variants across the LTR, gag, and pol, we compared CpG methylation across the env regions of the two subtypes, demonstrating that CpG methylation was comparable between the two at approximately 80% **(Figure 4d)**.

**Figure 4:**
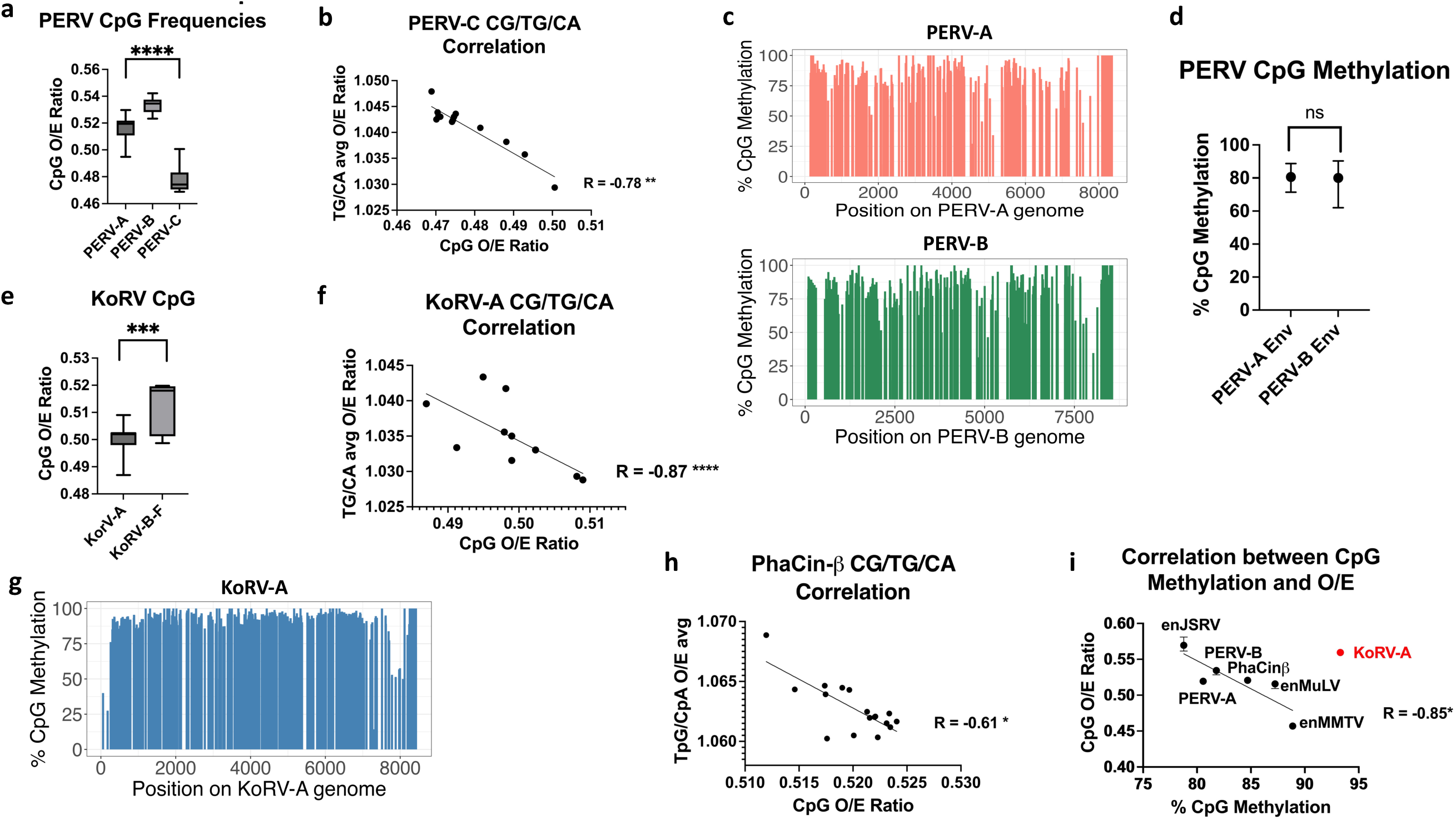
CpG content of seven vertebrate endogenous retroviruses corresponds with prior CpG methylation and transcriptional activity: **a)** Porcine Endogenous Retrovirus (PERV) variants exhibit significant variation in CpG content. **b)** PERV-C shows a significant negative correlation between CpG with TpG/CpA content, suggesting prior methylation. **c)** Bisulfite mapping with Bismark shows both PERV-A and PERV-B are heavily methylated across the entire provirus. **d)** PERV-A and PERV-B show no significant difference in median CpG methylation content. **e)** Endogenous Koala Retrovirus (KoRV-A) has significantly lower CpG content than the other KoRV subtypes. **f)** KoRV-A shows significant negative correlation between CpG and TpG/CpA frequency, suggesting impacts from DNA methylation. **g)** Bismark bisulfite mapping shows KoRV-A is heavily methylated across the entire provirus. **h)** Endogenous koala betaretrovirus phaCin-β shows a significant negative correlation between CpG and deamination products TpG/CpA. **i)** Median CpG % methylation plotted against median CpG observed/expected (O/E) ratios demonstrates a significant negative correlation between them for all endogenous retroviruses except KoRV-A (in red). Error bars in **(d)** represent interquartile ranges. P-values are reported for Kruskal-Wallis tests for multiple comparisons **(a)**, Mann-Whitney tests **(d,e)**, Spearman correlation **(b,f,h)**, and Pearson correlation **(i)**. *p < 0.05, **p < 0.01, *** p < 0.001, ****p < 0.0001

The koala gammaretrovirus, KoRV, is the youngest known endogenous retrovirus, as evidence points to it being in the midst of endogenizing. KoRV-A was the first subtype discovered and appears to be the most commonly endogenized variant, showing less association with disease than other subtypes. It is not entirely clear whether the other subtypes are endogenous or exogenous, but subtypes B-F are believed to be primarily exogenous.^9^ The CpG patterns of the other subtypes suggest they are less impacted by DNA methylation and potentially more transcriptionally active. For example, KoRV-A exhibits a significantly lower frequency of CpG dinucleotides than the other KoRV subtypes **(Figure 4e)**. Additionally, KoRV-A shows a significant correlation between CpG and TpG/CpA, once again suggesting DNA methylation plays a more significant role for this subtype **(Figure 4f)**, while no significant correlation was found between CpG O/E ratios and TpG/CpA ratios of the other subtypes (data not shown). Mapping of bisulfite converted koala DNA against the KoRV-A sequence showed heavy DNA methylation across the provirus in somatic liver tissue, which was also recently reported by ^29^ for testicular tissue **(Figure 4g)**, reinforcing the likely impact of DNA methylation upon the CpG composition of this provirus. Interestingly, the endogenous koala betaretrovirus, phaCin-β,^30^ also exhibited evidence for past DNA methylation, with a significant correlation between CpG O/E ratios with TpG/CpA ratios **(Figure 4h)**. Calculation of median %CpG methylation from mapping of bisulfite converted genomes for all endogenous retroviruses examined showed a significant correlation to median CpG O/E ratios, with KoRV-A proving to be a major outlier **(Figure 4i)**, perhaps reflecting its recent integration into the koala genome, as CpG frequencies have not yet decreased to the same extent as older endogenous retroviruses.

### Exogenous retroviruses exhibit more motifs associated with ’active’ histone modifications than endogenous variants

To evaluate differences in histone modifications across endogenous and exogenous retroviruses, we used Simple Enrichment Analysis (SEA) from the MEME suite^15^ to compare endogenous to exogenous proviral sequences and discover the most commonly enriched motifs in each. For this analysis we compared each of two types of endogenous MuLV sequences (modified polytropic (MPMV) and polytropic (PMV) to exogenous MuLV sequences. We also compared endogenous FeLV to exogenous FeLV-A, and endogenous JSRV to the two types of exogenous JSRV (Enzootic Nasal Tumor Virus, or ENTV, and Ovine Pulmonary Adenocarcinoma, or OPA).

Because mice have considerably more genome-wide histone modifications publicly available, a more thorough characterization of motifs associated with histone modifications across MuLVs was possible. We performed CpG island (CGI) analysis with Emboss Cpgplot^31^ to refine results for histone modifications typically confined to promoters and CGIs. CGI analysis showed that exogenous MuLV exhibited a CGI not present in the endogenous variants **(Figure 5a)**. Three statistically significant motifs for H3K27ac and H3K9ac, which are histone modifications associated with active transcription, were found in this CGI that were not located in the endogenous MuLVs **(Figure 5a, Supplementary Table S2)**. However, in PMV, which is more transcriptionally active than MPMV,^17^ a considerable number of motifs associated with both active (H3K9ac, H3K27ac, H3K4me3) and repressive (H3K27me3, H3K9me3) modifications were concentrated in the CGI traversing the U5 region of the 5’ LTR and the leader region **(Figure 5b)**. Interestingly, in addition to some of the motifs associated with ’active’ histone modifications, MPMV had multiple repressive H3K27me3 motifs throughout the provirus, including in the LTR, the CGI in the leader region, located in *gag*, and also in the CGI located at the *pol-env* junction **(Figure 5c)**.

**Figure 5:**
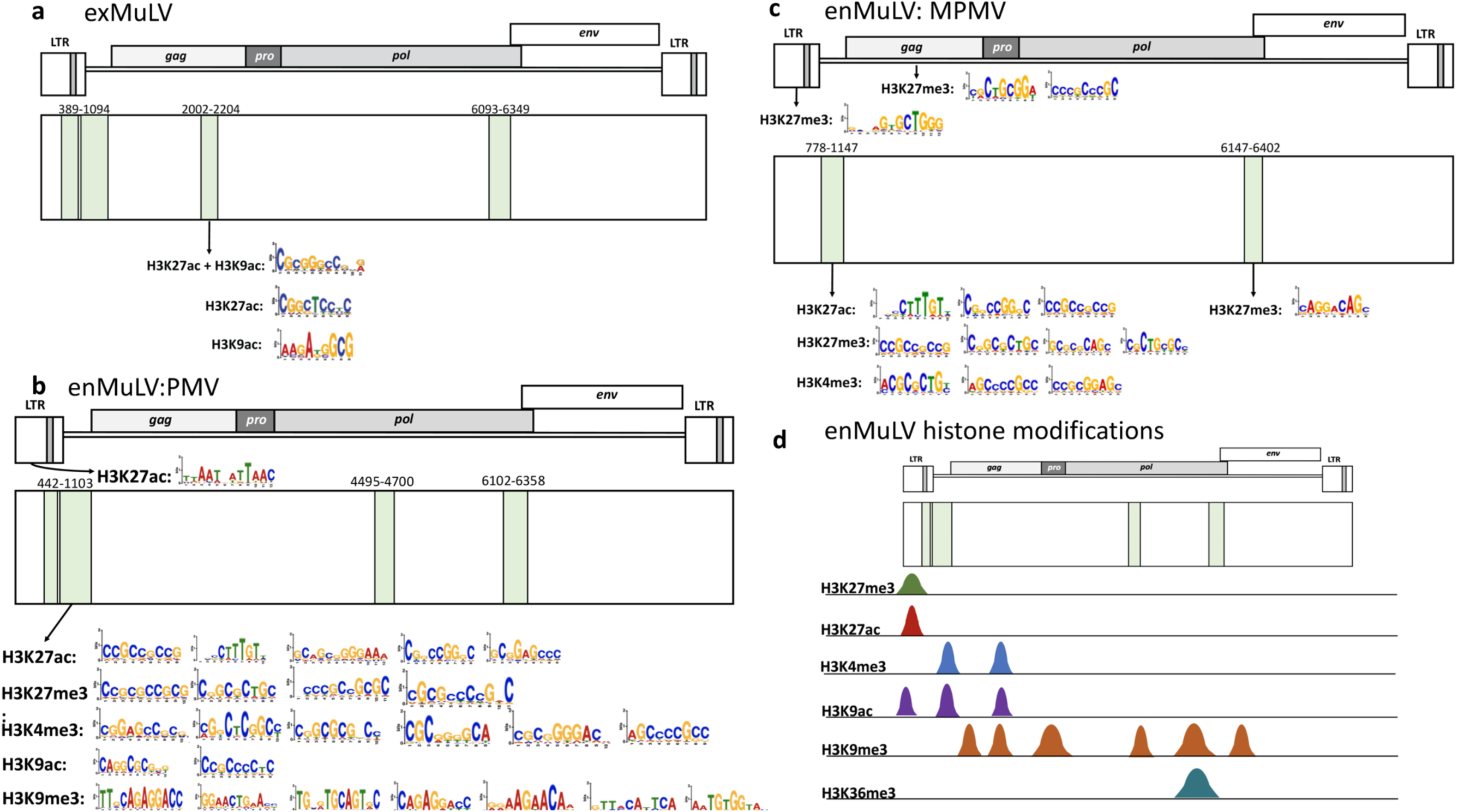
Murine leukemia virus (MuLV) shows motifs associated with histone modifications consistent with exogenous and variably active endogenous variants: **a)** Emboss Cpgplot and SEA analysis identify CpG islands (shaded in green) across the exogenous MuLV provirus, with one in particular containing H3K27ac and H3K9ac-associated motifs. **b)** Emboss Cpgplot and SEA analysis of polytropic endogenous MuLV (PMV) show heavy density of motifs associated with histone modifications within a CGI in the leader region, including those for active (H3K27ac, H3K4me3, H3K9ac) and repressive (H3K27me3 and H3K9me3) domains. **c)** Emboss Cpgplot and SEA analysis of endogenous modified polytropic endogenous MuLV (MPMV) shows widespread H3K27me3 motifs in the LTR and CGIs throughout the provirus. **d)** Visualization of ENCODE ChIP-Seq data from day 16 mouse embryos in the UCSC Genome Browser shows several histone modifications present in endogenous MuLV sequences (Figure details in **Table 3**). Further details on MuLV SEA analysis can be found in **Supplementary Table S2**.

**Table 3:**
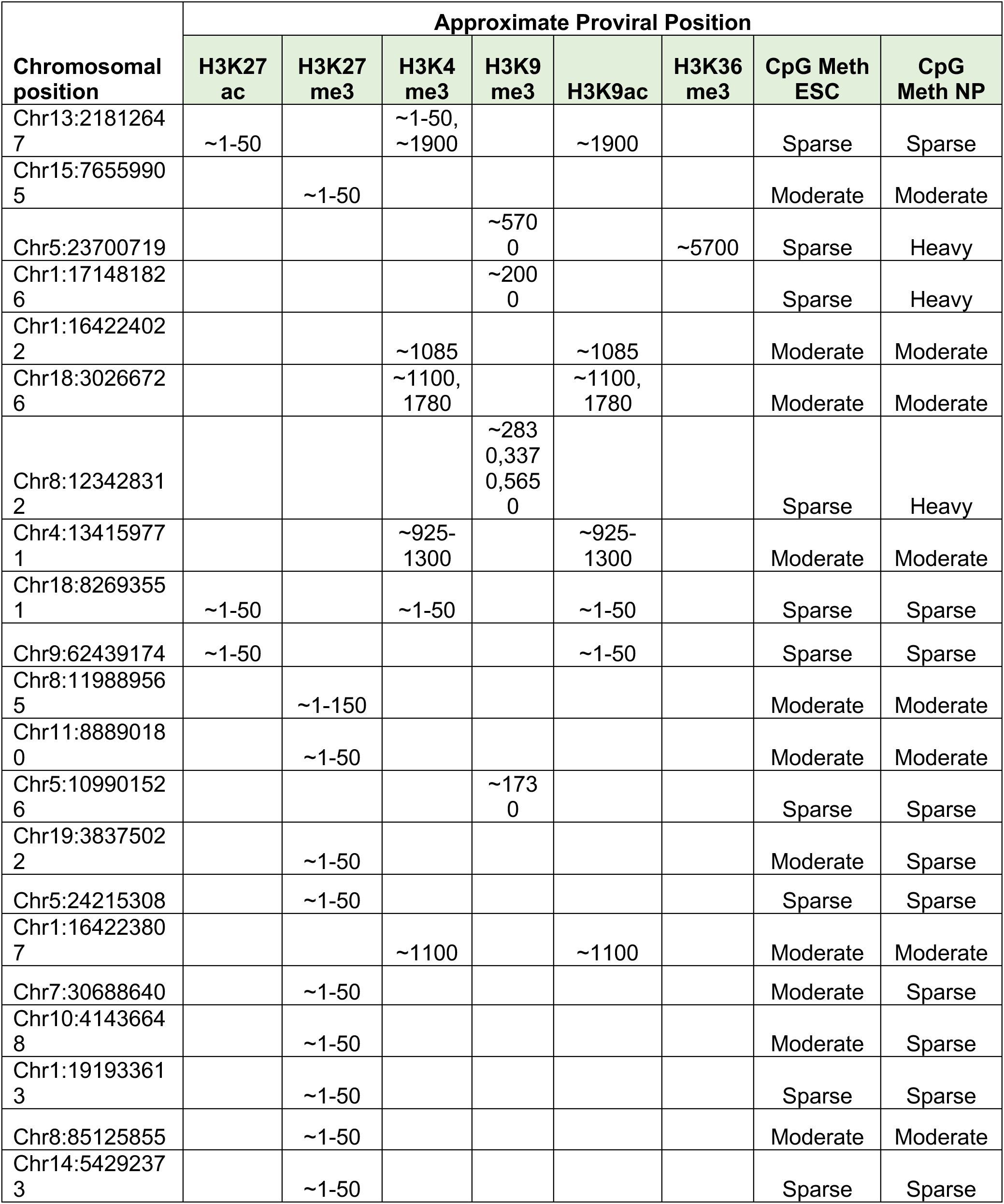
Histone modifications found in enMuLV using UCSC Genome Browser.

To attempt to validate these findings, we examined enMuLVs aligned with mouse genome assembly MM10 in the UCSC Genome Browser with ENCODE histone modification tracks. Surprisingly, 73% of enMuLV’s (57/78) showed no histone modifications at all, but a few did, summarized in **Table 3**. Repressive H3K9me3 and H3K27me3 were the most commonly found modifications, with some modifications associated with active transcription, including H3K27ac, H3K4me3, and H3K9ac **(Figure 5d)**. H3K27me3 and H3K27ac were most commonly located in the proximal LTR, while H3K4me3 and H3K9ac were distributed among CGIs and CpG-rich regions. H3K9me3 was broadly distributed across gene bodies, while H3K36me3 was located in the gene body of one provirus, with H3K9me3 marking the same spot. CpG methylation in embryonic stem cells and neural progenitors was usually found in sparse or moderate amounts in all proviruses examined and frequently overlapped histone modifications when they were present **(Table 3, Supplementary Figure S6a-b)**. However, mapping of bisulfite converted DNA from liver tissue showed heavy average CpG methylation across the provirus, suggesting CpG methylation might be more critical for proviral epigenetic regulation after differentiation **(Supplementary Figure S6c)**, although these data represent an average of proviral CpG methylation from different integrants at different sites vs. the individual sites examined for histone modifications.

To examine another gammaretrovirus, we generated H3K4me3 and H3K27ac motifs for the domestic cat and used SEA to assess enrichment throughout the endogenous and exogenous FeLVs. As both of these marks are associated with active transcription, there was a marked absence of any of these motifs meeting the relatively high significance threshold in enFeLVs (e-value < 0.05). However, exFeLV-A had multiple CGIs with which these modifications were associated **(Figure 6a, Supplementary Table S3)**.

**Figure 6:**
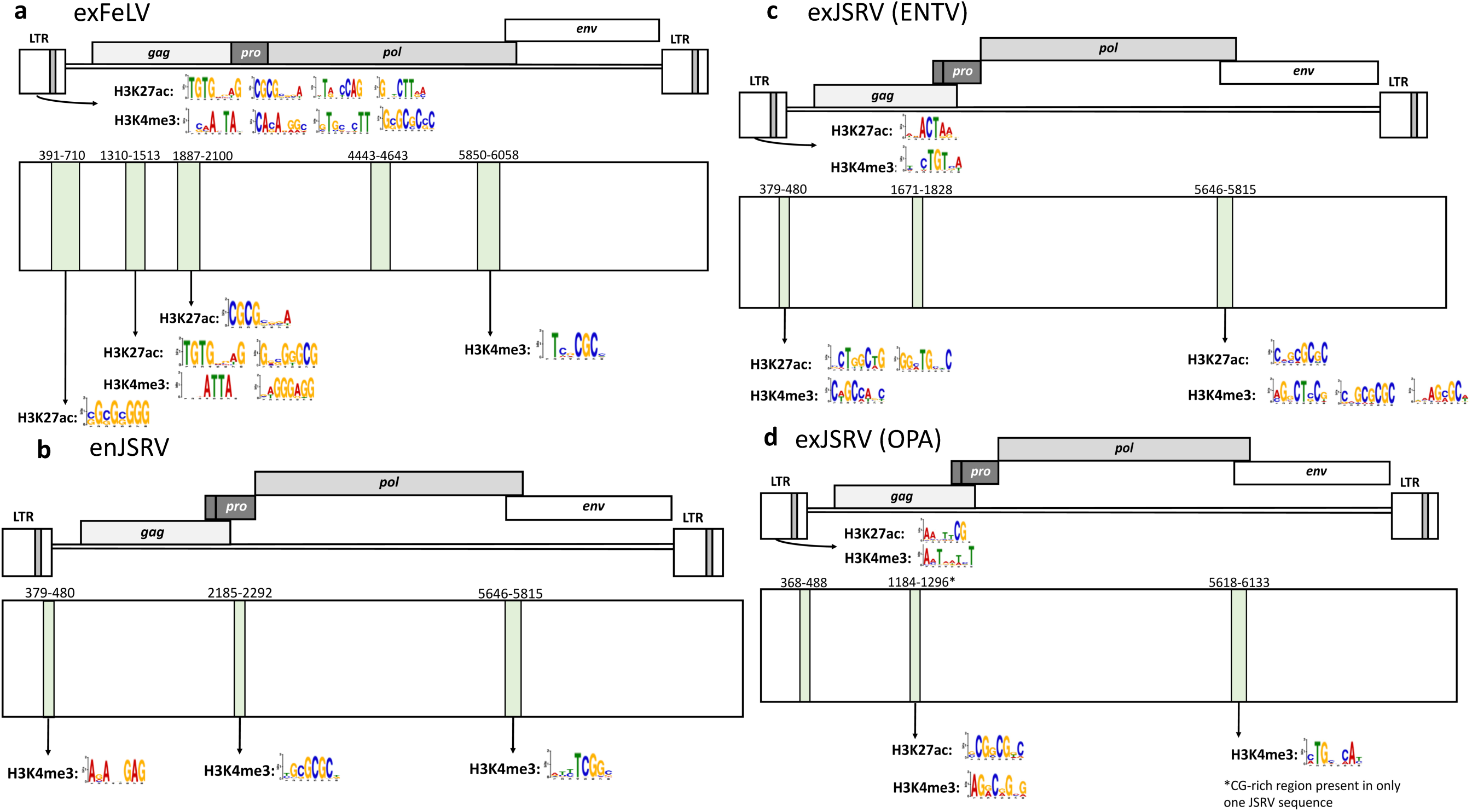
Exogenous Feline Leukemia Virus (FeLV) and Jaagsiekte Sheep Retrovirus (JSRV) variants contain more significant motifs associated with ’active’ histone modifications than endogenous variants. **a)** EMBOSS Cpgplot and SEA analysis of exogenous FeLV showed significant H3K27ac and H3K4me3 motifs in the LTR and CGIs throughout the provirus. Further details on SEA analysis can be found in **Supplementary Tables S3 and S4**. **b)** EMBOSS Cpgplot and SEA analysis show endogenous JSRV contains several CG-rich regions with only H3K4me3 motifs present and no H3K27ac. **c)** EMBOSS Cpgplot and SEA analysis of the exogenous Enzootic Nasal Tumor Virus (ENTV) JSRV variant show significant motifs present in the LTR, leader region, and *pol-env* junction for both H3K27ac and H3K4me3. **d)** EMBOSS Cpgplot and SEA analysis of the exogenous OPA JSRV variant shows several H3K27ac and H3K4me3-associated motifs within CG-rich regions primarily in the LTR and *gag*.

We also examined betaretroviruses for significantly enriched histone motifs. SEA analysis was overall unsuccessful for MMTV because too few full-length sequences were publicly available. However, JSRV analysis yielded some interesting findings. Both JSRV and MMTV exhibit considerably fewer conventional CGIs than the gammaretroviruses, so we narrowed the minimum length to 100 bp to pick up CpG-rich regions where histone modifying enzymes would be more likely to bind. All JSRV variants had short CpG stretches in the U5 region of the LTR and the CGI located in the pol-env junction. Endogenous JSRV had a CpG-rich region located in the *gag-pro-pol* junction, while the exogenous JSRVs had one more proximally located in *gag* **(Figure 6b-d)**. A few statistically significant H3K4me3 sites were found in the endogenous JSRV, but overall they were sparse **(Figure 6b, Supplementary Table S4)**. Both exogenous JSRVs had H3K27ac and H3K4me3 motifs in the proximal LTR **(Figure 6c-d)**. However, the ENTV JSRV variant had more of these motifs in the CpG-rich leader region and the *pol-env* junction, while the OPA variant had significant motifs in *gag,* suggesting these two exogenous variants are potentially regulated differently.

## Discussion

We have shown that beta- and gammaretroviruses frequently reflect the CpG content of their host promoters, but this is not always true in species where promoter CpG content is naturally lower. As shown in **Figure 1**, both beta- and gammaretroviruses can reflect host promoter CpG content, where in most species, ∼67% of promoters will contain CpG islands.^32^ Interestingly, promoter CpG content varies significantly between the different species examined **(Figure 1 and Supplementary Figure S2d)**, and those with lower CpG content appeared more divergent from the CpG content of their respective retroviruses. In some part, this could reflect retroviral evolution within other species first. KoRV in particular is hypothesized to have arisen in either rodents or bats based on phylogenomic studies.^33, 34^

JSRV is an older betaretrovirus thought to have integrated around 5-7 million years ago, but there is evidence of more recent endogenous variants having integrated only 200 years ago, and it is likely that the currently circulating exogenous variants continue to endogenize.^20^ The strong negative correlation between CpG and TpA/CpG motifs in the exogenous variants appears to reflect this evolutionary history, suggesting a long history of host CpG methylation and therefore repeated integration within the species. Overall, sheep promoters exhibit low CpG content compared to other species, resulting in a disparity between CpG content of promoters and JSRV. Originally, we hypothesized that the CpG content of the betaretroviral genomes might mirror that of host promoters due to a TSS integration bias. However, while this integration preference is not supported in the literature,^6^ the CpG content similarity to gammaretroviruses is still striking. The viral p12 protein of gammaretroviruses is known to tether the preintegration complex to nucleosomes via bromodomain and extraterminal domain (BET) proteins, playing a role in both integration and in nucleosome loading onto the provirus while targeting the PIC to CpG promoters and enhancers.^35, 36^ However, a tethering mechanism still has not been elucidated for betaretroviruses,^37^ so it is more difficult to hypothesize how their evolution might be impacted by host CpG content. The similarities between the genera in this respect could reflect convergent pressures for these endogenizing viruses. Instead of reflecting integration propensities, perhaps CpG content reflects the proviral tendency to harness promoter-like epigenetic machinery for successful transcription.

We have also shown that endogenization clearly impacts CpG content and CpG trinucleotide motif patterns, and that these results are likely due to CpG methylation. Every endogenous retrovirus to which we mapped bisulfite sequencing exhibited >80% methylation across the entire provirus **(Figures 2 and 4, Supplementary Figure S6c)**. CpG methylation has long been known to silence endogenous retroviruses during embryonic development.^38^ However, CpG methylation within gene bodies is generally not silencing and instead associated with active transcription.^39^ From an epigenetic standpoint, this could suggest that the entire provirus is treated similarly to a silenced host promoter. That said, even highly transcribed endogenous retroviruses (e.g. PERV^28^) showed heavy CpG methylation in somatic cells **(Figure 4c)**, calling into question what this mechanism achieves beyond transcriptional silencing. However, bisulfite mapping shows an average of all integrants in different regions, and it is possible that only a few escape repression to produce viral particles, while the majority are silenced.

Almost across the board, endogenous retroviruses exhibited significantly lower CpG content than their exogenous counterparts **(Figures 2-4)**, with some showing evidence more refined to methylation of particular CpG trinucleotide motifs. In general, these did not appear to reflect the methylation patterns of specific trinucleotide motifs in the host species. Different DNA methyltransferases will target different CpG trinucleotide motifs more heavily,^40, 41^ and it is unknown whether these might change between species. The varying promoter methylation patterns of CpG trinucleotides between the different species seem to suggest this is the case. Retroviral methylation is often initiated during embryonic development by *de novo* methyltransferases DNMT3a and DNMT3b.^40–42^ This may not be reflective of the overall methylation pattern in somatic cells, which is normally maintained by DNMT1.^43^

Time from initial ERV endogenization did not appear to correlate well with CpG methylation. For example, MMTV was believed to have endogenized initially around 20 million years ago **(Table 1)**, but it exhibited the most methylation and least amount of CpG loss compared to the other endogenous viruses **(Figure 4i)**, which integrated into their respective genomes initially ranging from 1-7 million years ago **(Table 1)**. However, it is critical to keep in mind that the endogenous retroviruses with exogenous counterparts, such as MMTV, are continuing to produce new endogenous variants that are younger and perhaps more prone to CpG methylation, and their integration sites likely also play a role in methylation frequencies. Therefore, while the approach of using CpG frequencies to estimate time from initial endogenization might work for stable integrants, it cannot be used to generalize time from endogenization for viruses that continue to circulate and create new integrants.

Using enMuLV data, we found that histone modifications were generally sparse among endogenous retroviruses, suggesting that the seemingly ubiquitous DNA methylation we observed **(Figures 2 and 4, Supplementary Figure S6c)** was more likely important for endogenous retroviral epigenetic regulation. Of proviruses that did have modifications, those associated with both active (H3K27ac, H3K9ac, H3K4me3, H3K36me3) and repressive (H3K27me3, H3K9me3) chromatin could be found **(Figure 5d**, **Table 3)**. It is probable the proviral histone modification pattern is dictated by the integration site environment and less by the endogenous provirus itself, which requires further exploration. However, there did appear to be significant differences in motifs associated with active and repressive histone modifications when comparing endogenous and exogenous retroviruses **(Figures 5 and 6)**, suggesting histone modifications could be playing an important role early after endogenization, but future work will need to evaluate this hypothesis. Alterations to histone modifications frequently precede DNA methylation for vertebrate gene regulation,^44^ so newly endogenized proviruses might be treated similarly. However, previous work on the more ancient human endogenous retroviruses suggested that DNA methylation was an earlier modification, while histone modifications were found in older and more degenerate LTRs.^45^ The profound (∼93%) DNA methylation across the provirus of the recently integrated KoRV-A supports this hypothesis **(Figure 4g)**. Additionally, KoRV-A defied the clear relationship between proviral DNA methylation and CpG frequency observed for older endogenous proviruses **(Figure 4i)**, suggesting changes to the viral sequence associated with methylated cytosine deamination have thus far been minimal, and supporting the idea that DNA methylation is a common initial epigenetic method for regulation of recent integrants.

Interestingly, with the exception of MMTV, the amount of CpG methylation across endogenous proviruses appeared to correlate roughly to their ages, with lower amounts of methylation corresponding to older integrants **(Table 1**, **Figure 4i)**, further supporting this hypothesis.

We confined our analysis of enMuLV histone modifications to sequences over 4000 bp, selecting for more intact sequences, and the majority of these sequences (73%) contained no histone modifications. Of those that did, CpG methylation was frequently present overlapping both activation-associated and repressive histone marks in neural progenitors **(Table 3, Supplementary Figure S6a-b)**. CpG methylation in embryonic stem cells and neural progenitors was almost ubiquitously present, but generally either in sparse or moderate amounts. However, heavy average CpG methylation is present across the enMuLV provirus in mouse liver **(Supplementary Figure S6c)**, suggesting DNA methylation is dynamic across enMuLV during development and also suggesting marked regulation via CpG methylation in somatic tissue.

In conclusion, retroviral endogenization shapes the sequence landscape of the provirus due to host epigenetic factors. CpG methylation clearly impacts proviral CpG content over time, while motifs associated with histone modifications appear to reflect the transcriptional state of the provirus, whether endogenous or exogenous. Generally speaking, of the three retroviruses we evaluated for motifs associated with histone modifications, exogenous variants appeared to have more prominent activation-associated motifs (i.e. H3K27ac and H3K4me3) than their endogenous counterparts relative to repression-associated motifs **(Figures 5 and 6)**. Endogenous MuLVs, for which we had more well-defined motifs to examine, exhibited more significant motifs associated with repression than exogenous variants, although those classified as PMV also harbored a number of activation-associated motifs, consistent with their known elevated transcriptional activity compared to other enMuLVs.^46^ While no one clear pattern emerged to distinguish endogenous from exogenous variants, we are still able to show that endogenization impacts the proviral sequence landscape in generally predictable ways, despite our analyses traversing multiple endogenous integrants in what are likely a wide variety of chromatin environments from different species.

## Methods

### CG content and trinucleotide motif O/E calculation

To calculate trinucleotide D-ratios (Observed/Expected or O/E ratios), a first order Markov model of conditional probabilities was used.^14, 47, 48^ The expected frequency of each motif is estimated using the individual base components with the following example equation:

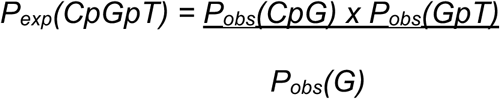

The D-ratio was calculated by dividing the P_obs_ of each trinucleotide by the P_exp_.

CG trinucleotide D-ratios were calculated for all proviral sequences using a Kmer tool previously published and provided by Dr. Diako Ebrahimi,^14^ as well as genomic sequences retrieved from each respective species using the *Ensembl* Biomart Tool.^49^ Genomic sequences extracted included 500 bp, 1 kb upstream, and 2 kb upstream of each TSS, as well as complete gene bodies and 1kb downstream of all Refseq annotated genes.

To identify and remove long terminal repeats from analyzed promoter sequences, we used LTRharvest^23^ from GenomeTools to identify and remove LTR sequences from all promoter regions in each species. Identification parameters included a minimum length of 10 bp, maximum length of 1000 bp, a minimum interval between LTRs of 15 bp with a maximum interval of 1000 bp.

### Bisulfite sequencing alignment and cytosine methylation calculation

Bisulfite converted genomes used in these analyses are reported in **Table 4**. SRA files obtained from NCBI were converted to fastq using the NCBI prefetch utility from the SRA toolkit. Fastq files were subjected to Trim Galore!.^50^ Alignment of retrieved promoter and retroviral sequences (accessions in **Table 5**) to bisulfite converted genomes was performed using Bismark software.^51^ Deduplication and methylation extraction were also performed using Bismark. CpG methylation reports generated from Bismark were used to calculate CpG methylation for CGA, CGC, CGG, and CGT trinucleotides.

**Table 4:** Bisulfite converted genomes from the NCBI <u>BioProjects</u> database used for this study.

| Species | Tissue | Bioproject | Experiment | Accession numbers |  |  |  |  |  |
| --- | --- | --- | --- | --- | --- | --- | --- | --- | --- |
| Cat | Blood | PRJNA16726 | SRX548709 | SRR1293903 | SRR1293905 |  |  |  |  |
| Sheep | Skeletal Muscle | PRJNA622418 | SRX8073310 | SRR11497387 |  |  |  |  |  |
| Pig | Blood | PRJEB55066 | ERX11037418 | ERR11635337 |  |  |  |  |  |
| Koala | Kidney | PRJNA629340 | SRX8207656 | SRR11646494 | SRR11646495 |  |  |  |  |
| Mouse | Liver | PRJNA416505 | SRX3347189 | SRR6239777 | SRR6239784 | SRR6239792 | SRR6239799 | SRR6239807 | SRR6239814 |
|  |  |  |  | SRR6239778 | SRR6239785 | SRR6239793 | SRR6239801 | SRR6239808 | SRR6239815 |
|  |  |  |  | SRR6239779 | SRR6239786 | SRR6239794 | SRR6239802 | SRR6239809 | SRR6239816 |
|  |  |  |  | SRR6239780 | SRR6239787 | SRR6239795 | SRR6239803 | SRR6239810 |  |
|  |  |  |  | SRR6239781 | SRR6239788 | SRR6239796 | SRR6239804 | SRR6239811 |  |
|  |  |  |  | SRR6239782 | SRR6239789 | SRR6239797 | SRR6239805 | SRR6239812 |  |
|  |  |  |  | SRR6239783 | SRR6239790 | SRR6239798 | SRR6239806 | SRR6239813 |  |

**Table 5:** Sources and NCBI <u>GenBank</u> Accession numbers for proviral sequences used in this study.

| Gammaretroviruses |  |  |  |  |  |
| --- | --- | --- | --- | --- | --- |
| *enMuLV*<sup>25</sup> |  |  |  |  |  |
| Mpmv1 | Mpmv6 | Pmv14 | Pmv22 | Xmv10 | Xmv42 |
| Mpmv10 | Mpmv7 | Pmv15 | Pmv23 | Xmv12 | Xmv43 |
| Mpmv11 | Mpmv8 | Pmv16 | Pmv24 | Xmv13 | Xmv8 |
| Mpmv12 | Mpmv9 | Pmv17 | Pmv4 | Xmv15 | Xmv9 |
| Mpmv13 | Pmv1 | Pmv18 | Pmv5 | Xmv16 |  |
| Mpmv2 | Pmv10 | Pmv19 | Pmv6 | Xmv17 |  |
| Mpmv3 | Pmv11 | Pmv2 | Pmv7 | Xmv18 |  |
| Mpmv4 | Pmv12 | Pmv20 | Pmv8 | Xmv19 |  |
| Mpmv5 | Pmv13 | Pmv21 | Pmv9 | Xmv41 |  |
| **exMuLV** |  |  |  |  |  |
| KY574504.1 | KY574507.1 | KY574510.1 | KY574513.1 | KY574516.1 | MG525170.1 |
| KY574505.1 | KY574508.1 | KY574511.1 | KY574514.1 | KY574517.1 |  |
| KY574506.1 | KY574509.1 | KY574512.1 | KY574515.1 | KY574518.1 |  |
| **enFeLV** |  |  |  |  |  |
| AY364318.1 | LC196053.1 | LC196055.1 | OP595707.1 |  |  |
| AY364319.1 | LC196054.1 | OP595706.1 | OP595708.1 |  |  |
| **exFeLV-A** |  |  |  |  |  |
| AB672612.1 | AB672612.1 | M18247.1 | MZ964580.1 |  |  |
| AF052723.1 | AF052723.1 | MT129531.1 | NC_001940.1 |  |  |
| **KoRV-A full-length** |  |  |  |  |  |
| AB721500.1 | KF786280.1 | KF786282.1 | KF786284.1 | KF786286.1 |  |
| KC779547.1 | KF786281.1 | KF786283.1 | KF786285.1 | NC_039228.1 |  |
| **KoRV-A env** |  |  |  |  |  |
| AB721500.1 | JQ244835.1 | JQ244838.1 | KF786281.1 | KF786284.1 | NC_039228.1 |
| AB823238.1 | JQ244836.1 | JQ244839.1 | KF786282.1 | KF786285.1 |  |
| AF151794.2 | JQ244837.1 | KF786280.1 | KF786283.1 | KF786286.1 |  |
| **KoRV-B-F env** |  |  |  |  |  |
| AB822553.1 | AB828005.1 | KP792564.1 | KU533852.1 |  |  |
| AB828004.1 | KC779547.1 | KP792565.1 | KU533853.1 |  |  |
| **PERV-A** |  |  |  |  |  |
| AF435966.1 | AY099323.1 | HM159246.1 | KY484771.1 |  |  |
| AJ133817.1 | EF133960.1 | HQ540591.1 | OR574834.1 |  |  |
| **PERV-B** |  |  |  |  |  |
| AJ133816.1 | AJ293657.1 | EU523109.1 | HQ536010.1 | HQ540594.1 |  |
| AJ133818.1 | AY056035.1 | HQ536008.1 | HQ536011.1 | HQ540595.1 |  |
| AJ279057.1 | AY099324.1 | HQ536009.1 | HQ540593.1 |  |  |
| **PERV-C** |  |  |  |  |  |
| AF038599.1 | AY570980.1 | HQ536014.1 | KC116219.1 | KY352351.2 |  |
| AF038600.1 | HQ536012.1 | HQ536015.1 | KC116220.1 | MT984604.1 |  |
| AM229312.2 | HQ536013.1 | HQ536016.1 | KC116221.1 |  |  |
| **Betaretroviruses** |  |  |  |  |  |
| **enMMTV** |  |  |  |  |  |
| AC122322.4 | AF228550.1 | NG_005612.1 |  |  |  |
| AC140344.3 | AL683884.26 |  |  |  |  |
| **exMMTV** |  |  |  |  |  |
| AF033807.1 | AF228552.1 | M15122.1 |  |  |  |
| AF228551.1 | D16249.1 | NC_001503.1 |  |  |  |
| **enJSRV** |  |  |  |  |  |
| AF136224.1 | EF680297.1 | EF680302.1 | EF680307.1 | EF680314.1 | MF175071.1 |
| AF136225.1 | EF680298.1 | EF680303.1 | EF680308.1 | MF175067.1 |  |
| AF153615.1 | EF680299.1 | EF680304.1 | EF680309.1 | MF175068.1 |  |
| DQ838493.1 | EF680300.1 | EF680305.1 | EF680310.1 | MF175069.1 |  |
| EF680296.1 | EF680301.1 | EF680306.1 | EF680312.1 | MF175070.1 |  |
| **exJSRV OPA** |  |  |  |  |  |
| AF105220.1 | CQ964469.1 | KP691837.1 | NC_001494.1 |  |  |
| AF357971.1 | DQ838494.1 | M80216.1 |  |  |  |
| **exJSRV ENTV** |  |  |  |  |  |
| FJ744146.1 | FJ744149.1 | GU292315.1 | GU292318.1 | PP707036.1 |  |
| FJ744147.1 | FJ744150.1 | GU292316.1 | KC189895.1 |  |  |
| FJ744148.1 | GU292314.1 | GU292317.1 | NC_007015.1 |  |  |
| **phaCin-β<sup>30</sup>** |  |  |  |  |  |
| pCi1002 | pCi237 | pCi366 | pCi47 | pCi921 | pCi989 |
| pCi194 | pCi273 | pCi414 | pCi643 | pCi933 | pCi998 |
| pCi233 | pCi303 | pCi457 | pCi786 | pCi983 |  |

### Histone motif generation

Motifs for all mouse histone modifications were retrieved from.^52^ Epigram was used as described in ^52^ to generate motifs for H3K4me3 and H3K27ac in sheep and domestic cats using the following data sets: H3K4me3 and H3K27ac from sheep liver GSE206736^21^ aligned with the Ovis aries genome assembly Oar_rambouillet_v1.0, and H3K4me3 and H3K27ac from cat heart GSE182952 aligned with the *Felis catus* genome assembly F.catus_Fca126_mat1.0.

### CpG Island Identification

For CpG Island identification, consensus sequences for each provirus were generated using MacVector® software. Sequences were entered into Emboss Cpgplot^31^ using a window size of 100, a minimum length of 200, a minimum observed CpG frequency of 0.6, and a minimum CpG percentage of 50. For JSRV sequences, CGI minimum lengths were dropped to 100.

### SEA analysis

For Simple Enrichment Analysis (SEA), we used the SEA tool from the MEME suite.^15^ Histone modification motif analyses for exogenous retrovirus variants were submitted for enrichment compared to their corresponding endogenous variants as negative controls, and vice versa with histone motifs generated from Epigram analyzed for enrichment as described above with an E-value cutoff of 0.05. Motifs for H3K4me3, H3K27ac, and H3K27me3 were retrieved only if they aligned with CpG islands from the corresponding proviruses being analyzed. In all cases, only sequence motifs enriched in all inputted sequences with no more than one of the negative control sequences, or one less than all inputted sequences with no negative control sequences, were considered significant, even when E-value and CGI criteria were met.

### enMuLV histone modification identification

Consensus sequences for endogenous MuLV variants MPMV, PMV, and XMV were all entered into the UCSC Genome Browser BLAT tool.^53, 54^ ENCODE histone tracks for H3K27ac, H3K27me3, H3K9ac, H3K9me3, H3K4me3, H3K36me3, and H3K4me1 from forebrain, liver, kidney, and intestine from day 16 embryonic mice aligning ζ 4000 bp of enMuLV sequence were visualized to identify sites of modification throughout the provirus. ENCODE tracks for CpG methylation in mouse embryonic stem cells and neural progenitor cells were also included.

### Statistical analysis

Statistical analyses and data visualization were performed using GraphPad Prism® software version 10.4.2 for macOS, as well as the stats and rstatix packages in R for statistics and ggplot2 for visualization. Multiple comparisons for CpG motifs between proviruses and promoters were performed using the Kruskal-Wallis test for multiple comparisons followed by Dunn’s tests. Two-sided independent t-tests or Mann-Whitney tests were used for individual comparisons where appropriate. All correlation analyses were performed using Spearman’s rank correlation test with a 95% confidence interval. Pearson correlation was used in one case **(Figure 4i)** after the normality assumption was confirmed by Shapiro-Wilk and Komogorov-Smirnov tests. In all analyses, statistical significance was defined as p-value < 0.05.

### Phylogenetic analysis

Phylogenetic analysis for JSRV sequences reported in **Supplementary Figure S2** were conducted using MacVector® software. Neighbor-joining trees were built using 1000 bootstraps from from enJSRV and exJSRV sequences aligned with Clustal W.

## Data Availability Statement

### Accession numbers

Chip-seq data from <u>GSE206736</u> was retrieved to analyze for H3K4me3 and H3K27ac motifs in sheep. Chip-seq data from <u>GSE182952</u> was retrieved to analyze for H3K4me3 and H3K27ac motifs in domestic cats.

**Table 4** lists bisulfite converted genome sequences accessed on the NCBI BioProjects database with the following Project, SRA Experiment, and Accession numbers:

**Table 5** lists accession numbers for all viral sequences accessed on NCBI Genbank:

### Tools and software

Genomic sequence retrieval was performed with <u>Ensembl Biomart</u>.

<u>LTRHarvest</u>^23^ from <u>GenomeTools</u> was used to extract LTR sequences from promoters.

SRA files obtained from NCBI were converted to fastq using the NCBI prefetch utility from the <u>SRA toolkit</u>. Fastq files were trimmed using <u>Trim Galore!</u>.^50^ Bisulfite converted sequences were aligned with <u>Bismark</u>^51^ software.

<u>Epigram</u>^52, 55^ was used to predict motifs associated with histone modifications.

Proviral alignments, consensus generation, and phylogenetic analysis was performed with <u>MacVector®</u> software.

<u>Emboss Cpgplot</u>^31^ was used for CpG Island identification.

The <u>SEA</u> tool from the <u>MEME</u> suite^15^ was used for Simple Enrichment Analysis.

The <u>UCSC Genome Browser BLAT</u> tool^53, 54^ was used to locate endogenous proviral sequences in the mouse genome for histone modification identification with <u>ENCODE histone modification tracks</u>.

Statistical analyses were performed using <u>GraphPad PrismÒ</u> software, as well as the <u>rtstatix</u> package in <u>R</u>. Visualization was also performed with <u>GraphPad PrismÒ</u> software and <u>ggplot2</u> in R.

## Acknowledgements

This work was supported by the James B. Pendleton Charitable Trust, UC San Diego faculty research funds, the California HIV Research Program CHRP H24BD7869 (LaMere), and grants from the National Institutes of Health: 1K01OD026565 (LaMere), R01NS137852 (LaMere), and R01HG009626 (Wang). We would also like to acknowledge Dr. Diako Ebrahimi for the provision of the Python code used for the Kmer analysis in this manuscript.

