## Supplementary Material for "Epigenetic motifs distinguishing endogenous from exogenous retroviral integrants"

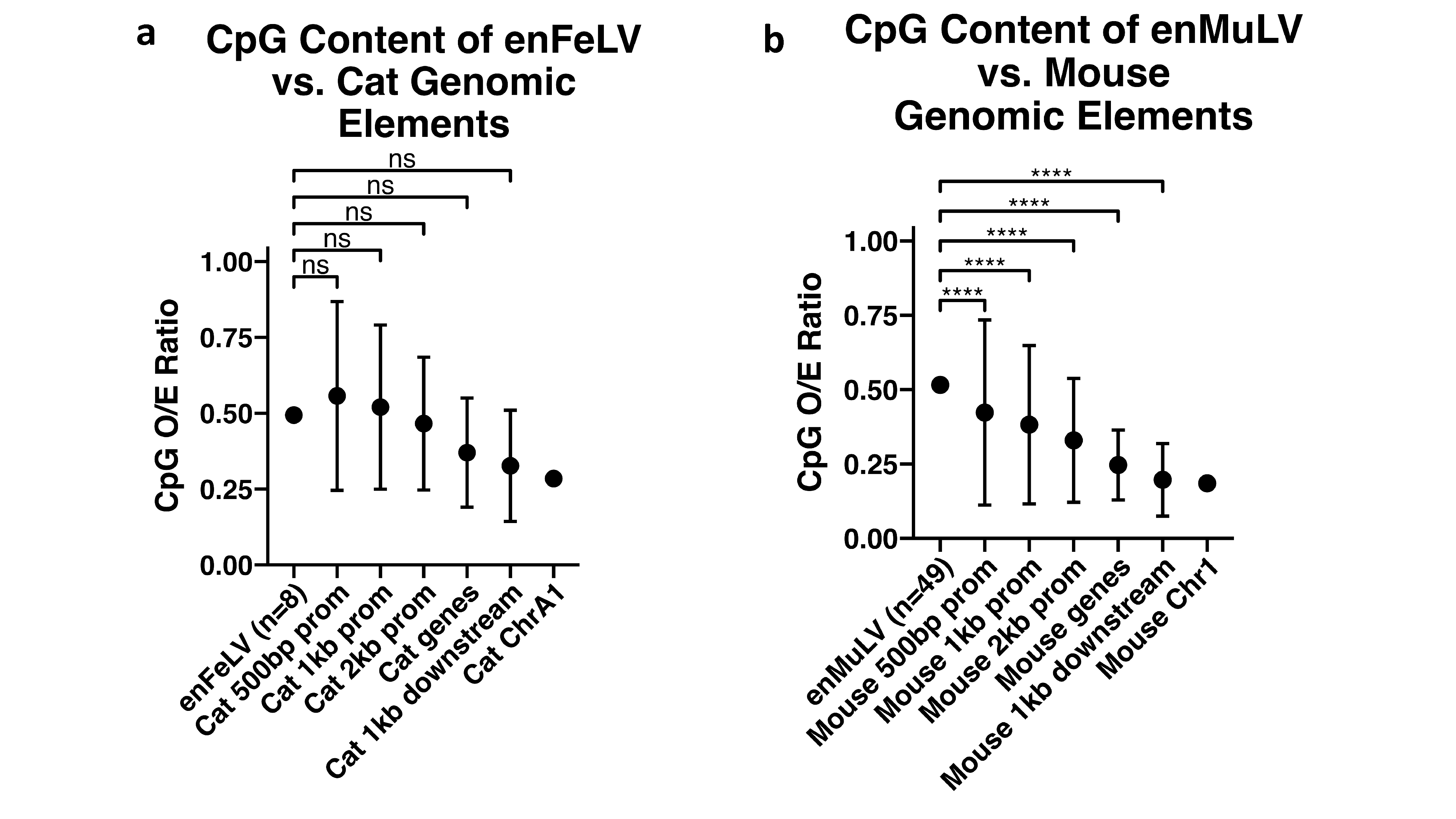

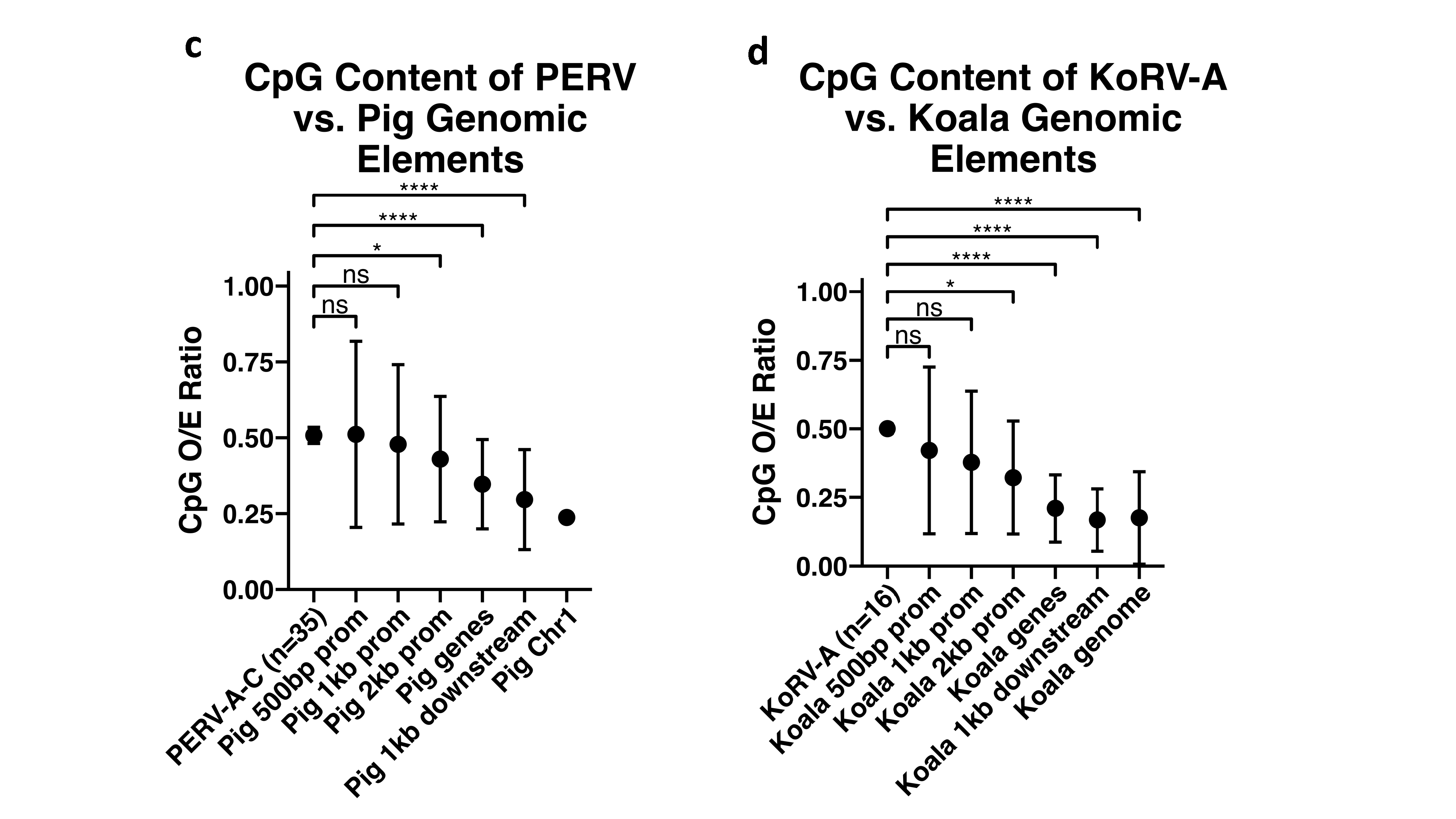

**Supplementary Figure S1:** Mean CpG content of endogenous gammaretroviruses vs. promoters (i.e. 500 bp upstream of the TSS, 1 kb upstream of the TSS, 2 kb upstream of the TSS), gene bodies, 1 kb downstream of genes, and entire chromosomes (or genome in the case of koala). **a)** Endogenous FeLV vs. cat genomic elements **b)** endogenous MuLV vs. mouse genomic elements **c)** Porcine Endogenous Retroviruses A-C vs. pig genomic elements **d)** KoRV-A vs. koala genomic elements. P-values are reported for the Kruskal-Wallis test for multiple comparisons with Dunn's test, while error bars represent interquartile ranges. *p < 0.05, **p < 0.01, ***p < 0.001, ****p < 0.0001

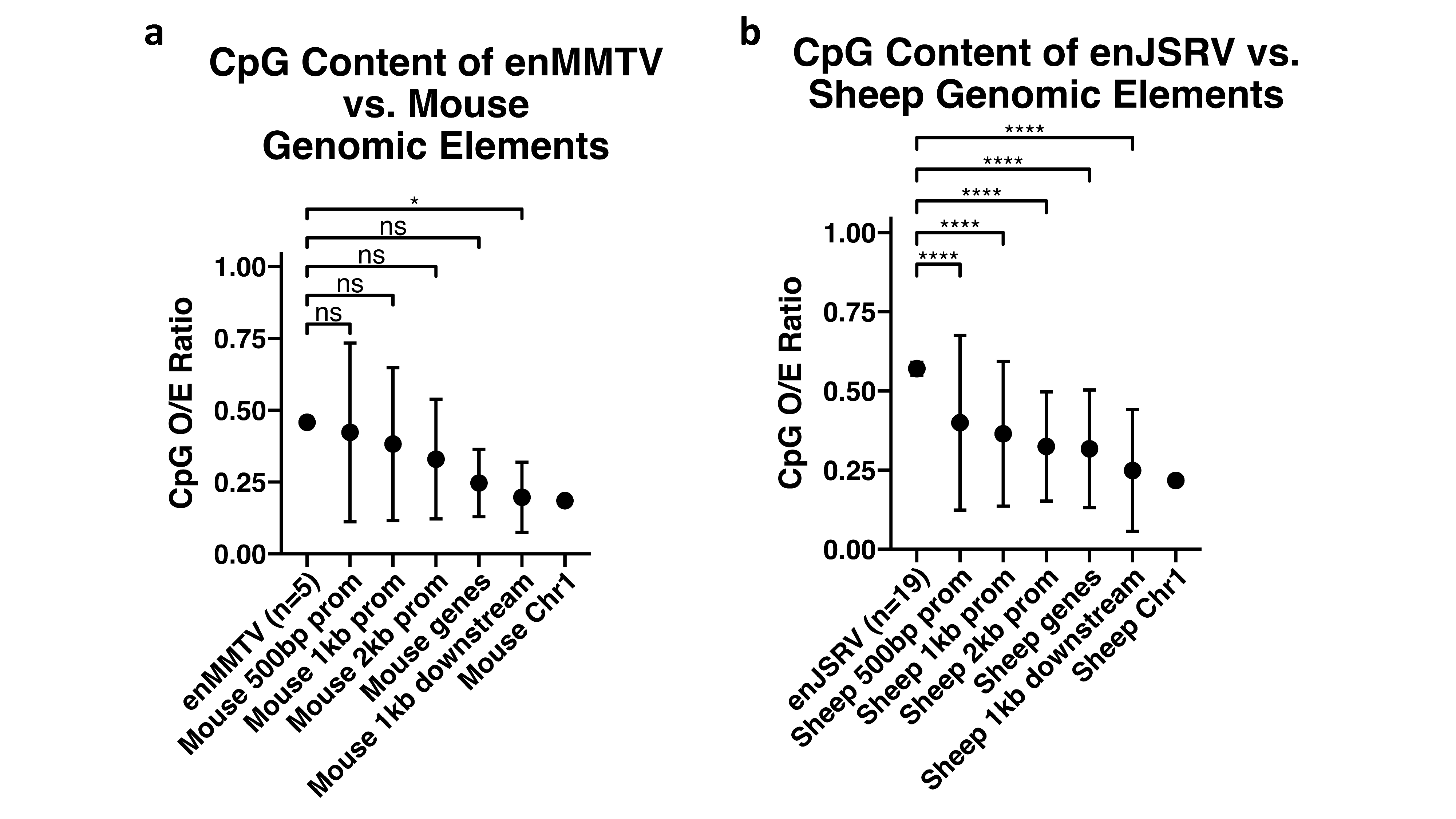

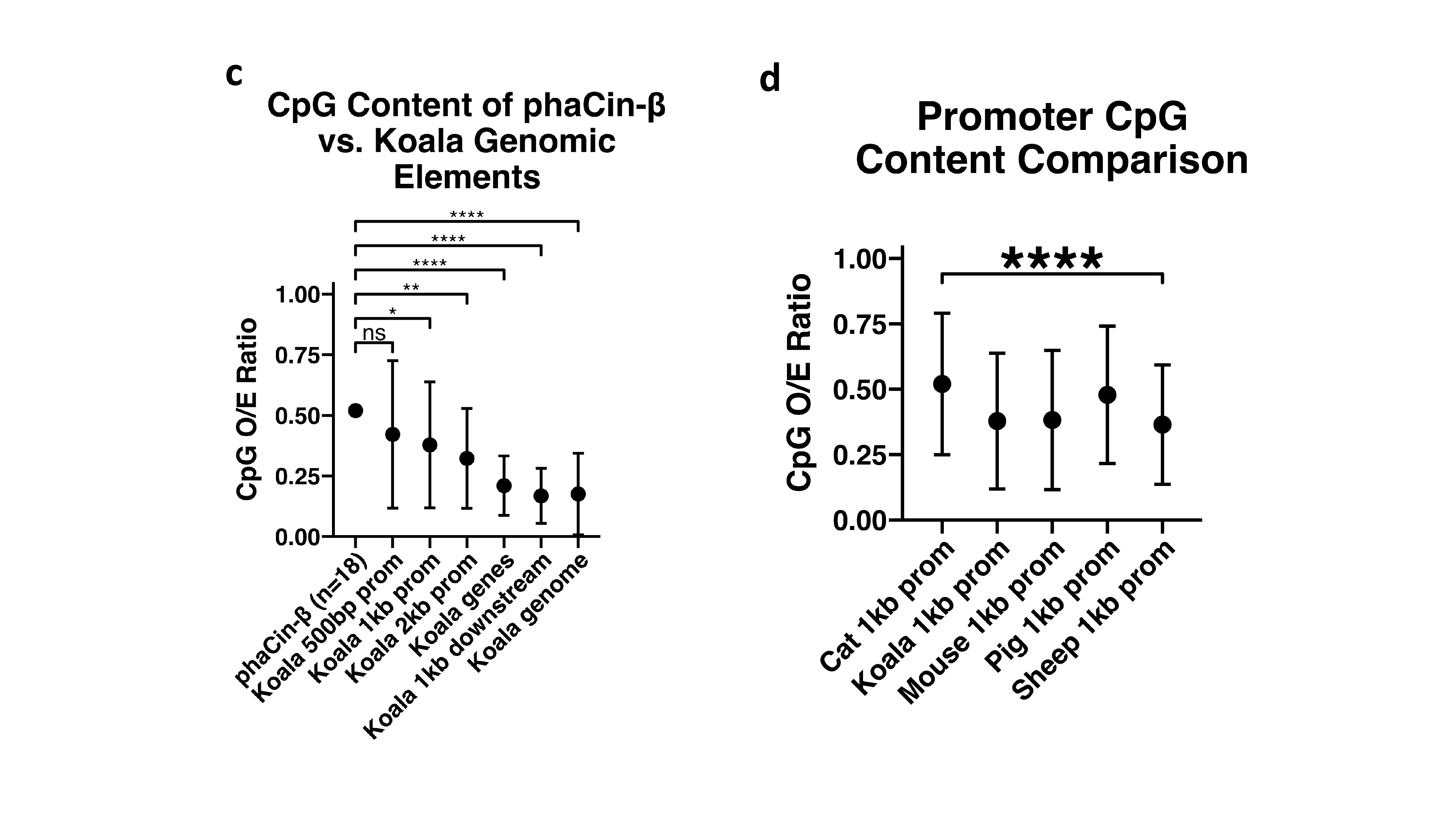

**Supplementary Figure S2:** CpG content of endogenous betaretroviruses vs. promoters (i.e. 500 bp upstream of the TSS, 1 kb upstream of the TSS, 2 kb upstream of the TSS), gene bodies, 1 kb downstream of genes, and entire chromosomes (or genome in the case of koala). **a)** Endogenous MMTV vs. mouse genomic elements **b)** endogenous JSRV vs. sheep genomic elements **c)** phaCin-β vs. koala genomic elements **d)** Comparison of promoter (1 kb upstream of TSS) CpG content for all 5 species. P-values for **a-c** are reported for the Kruskal-Wallis test for multiple comparisons with Dunn's test, while p-value for **d** is for Kruskal-Wallis test alone. Error bars represent interquartile ranges. *p < 0.05, **p < 0.01, ***p < 0.001, ****p < 0.0001

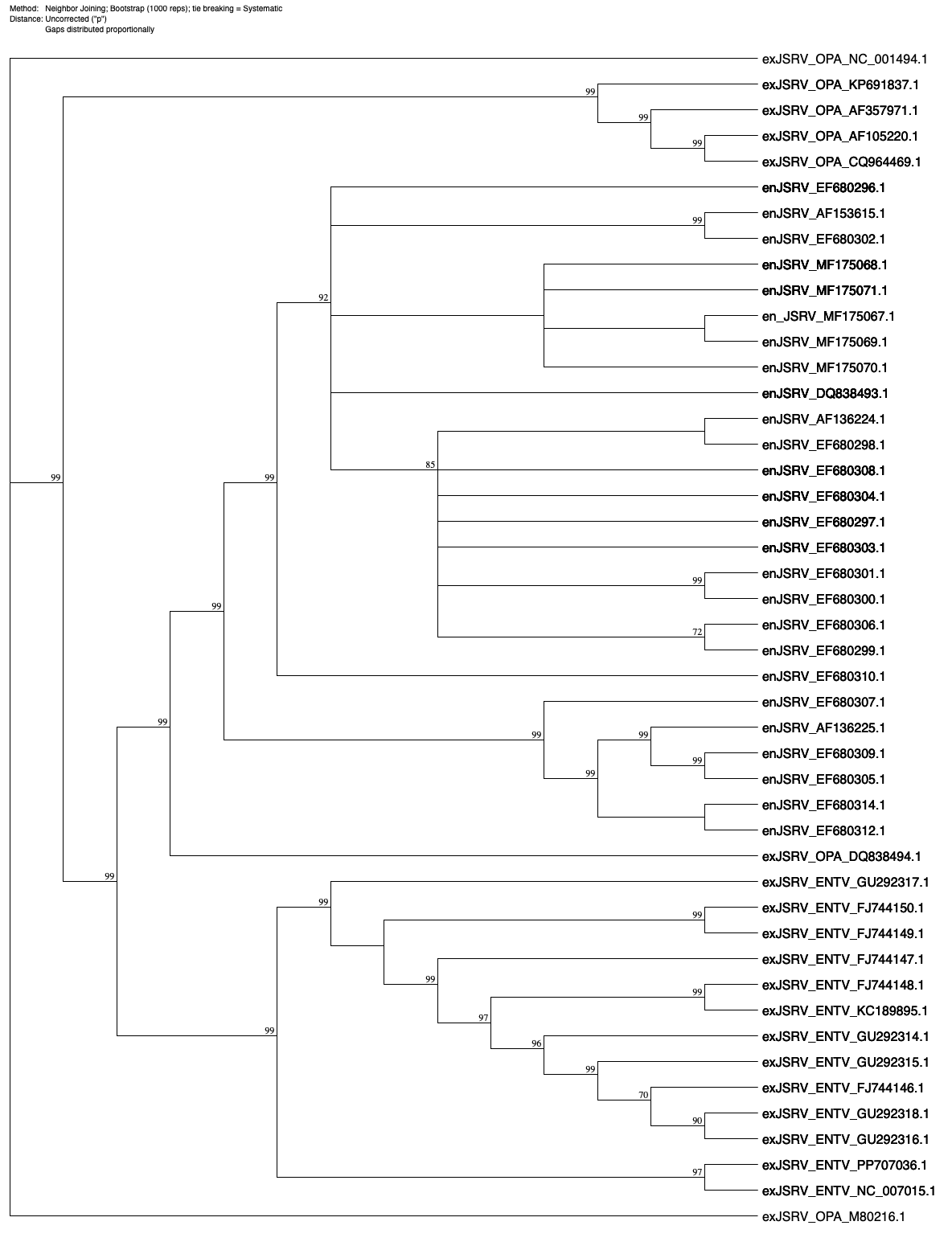

**Supplementary Figure S3:** Neighbor joining phylogenetic tree of endogenous JSRV (enJSRV), exogenous JSRV of Enzootic Nasal Tumor Virus (exJSRV_ENTV) and exogenous JSRV of ovine Pulmonary Adenocarcinoma (exJSRV_OPA). The tree was constructed using MacVector® software with 1000 bootstraps.

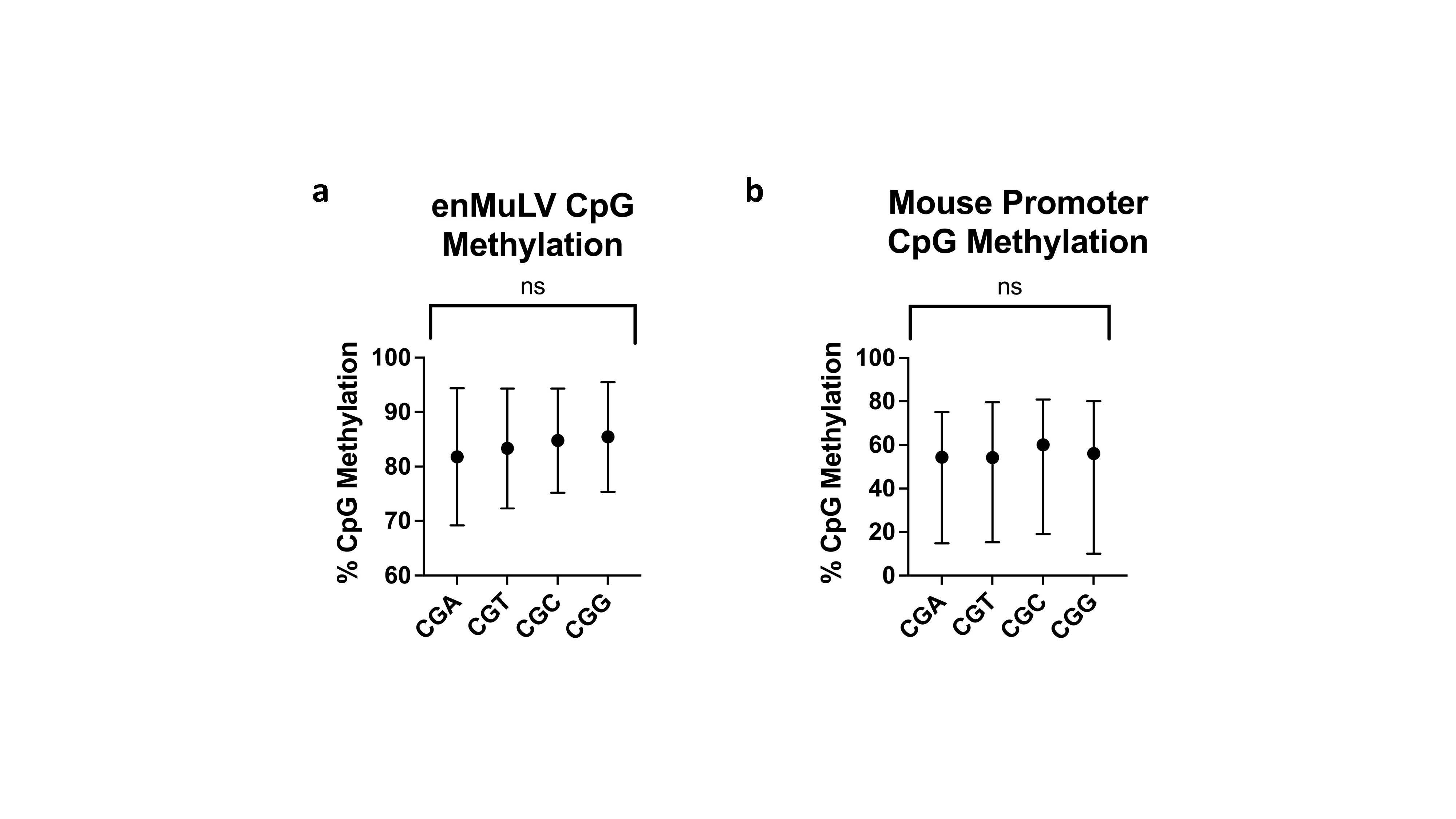

**Supplementary Figure S4:** CpG methylation in both endogenous MuLV and mouse promoters shows no preference for specific trinucleotide motifs. **a)** CpG methylation across CGA, CGT, CGC, and CGG motifs in endogenous MuLV. **b)** CpG methylation across CGA, CGT, CGC, and CGG motifs in mouse promoters. Medians with interquartile ranges represented. P-values are reported for Kruskal-Wallis test for multiple comparisons.

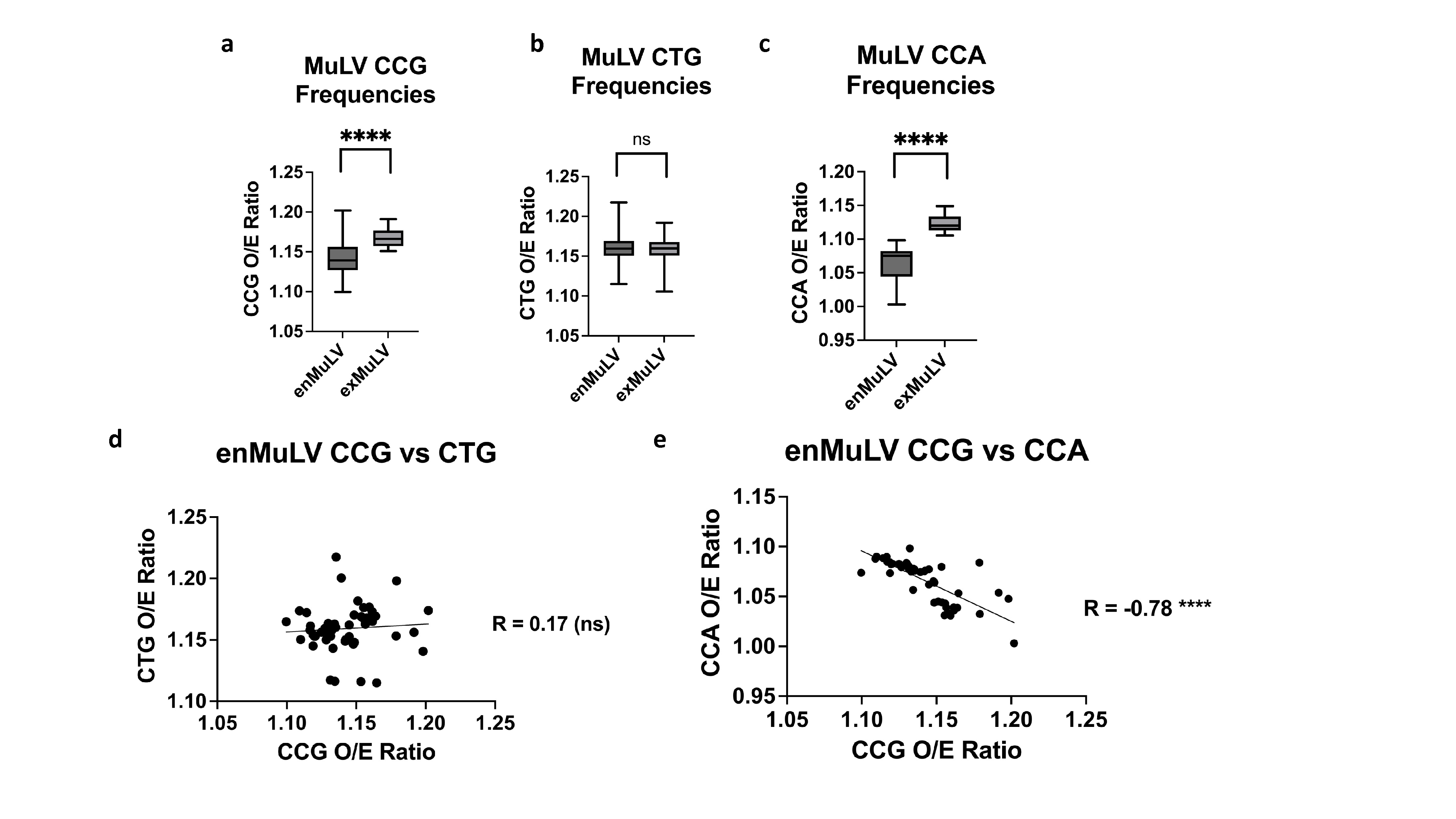

**Supplementary Figure S5:** Endogenous MuLV shows asymmetric CCG deamination. **a)** enMuLV exhibits significantly lower CCG than exMuLV. **b)** enMuLV shows no significant difference between CCG deamination product CTG with exMuLV. **c)** enMuLV exhibits a significantly lower CCA O/E (i.e. CCG deamination product) than exMuLV, contrary to expectations. **d)** No significant correlation is found between CCG and deamination product CTG in enMuLV. **e)** A strong negative correlation is found between CCG and deamination product CCA in enMuLV. P-values for a-c represent Mann-Whitney tests, while R-values and p-values for (d) and (e) represent Spearman correlation. ns p > 0.05, ****p < 0.0001

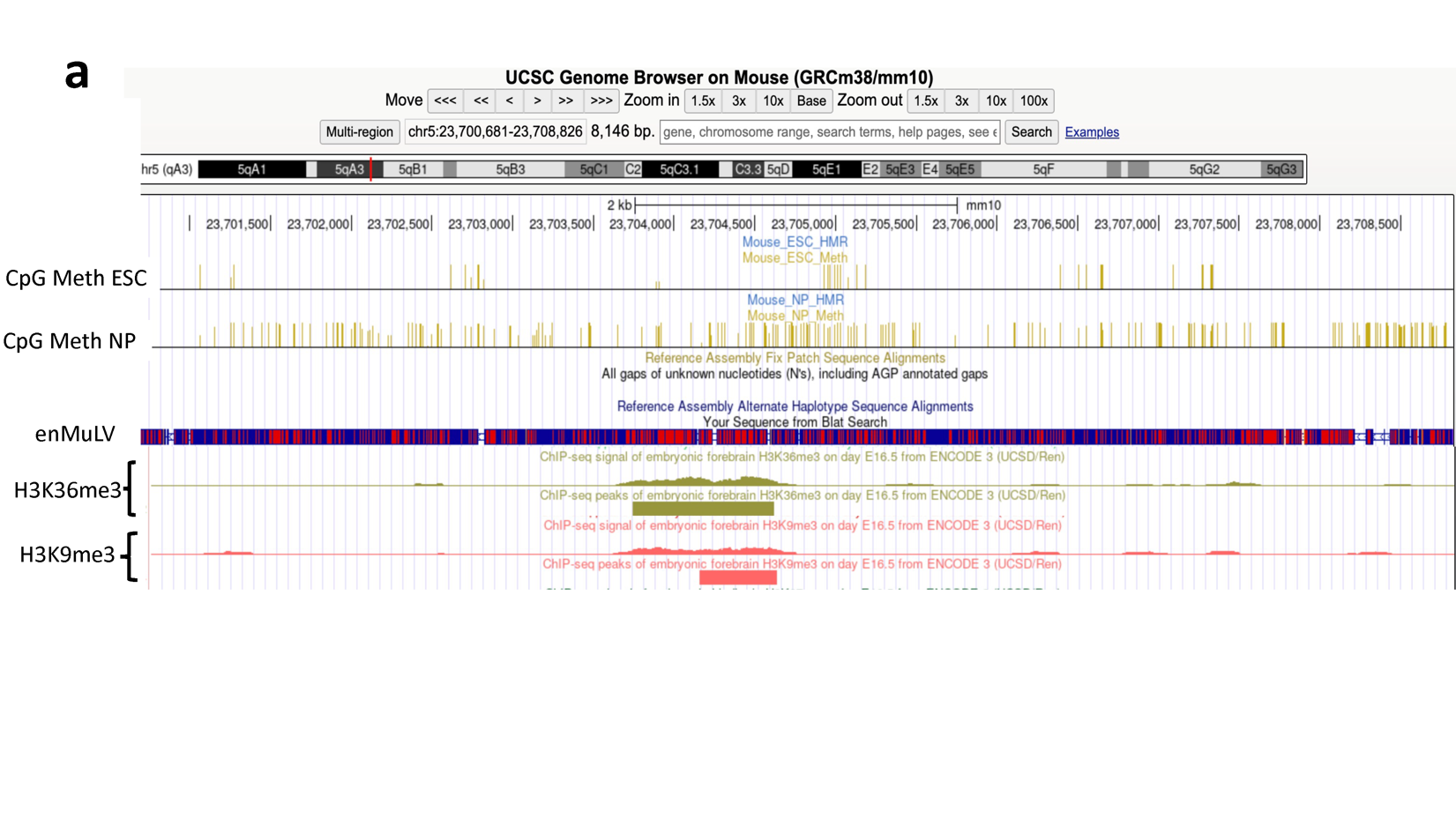

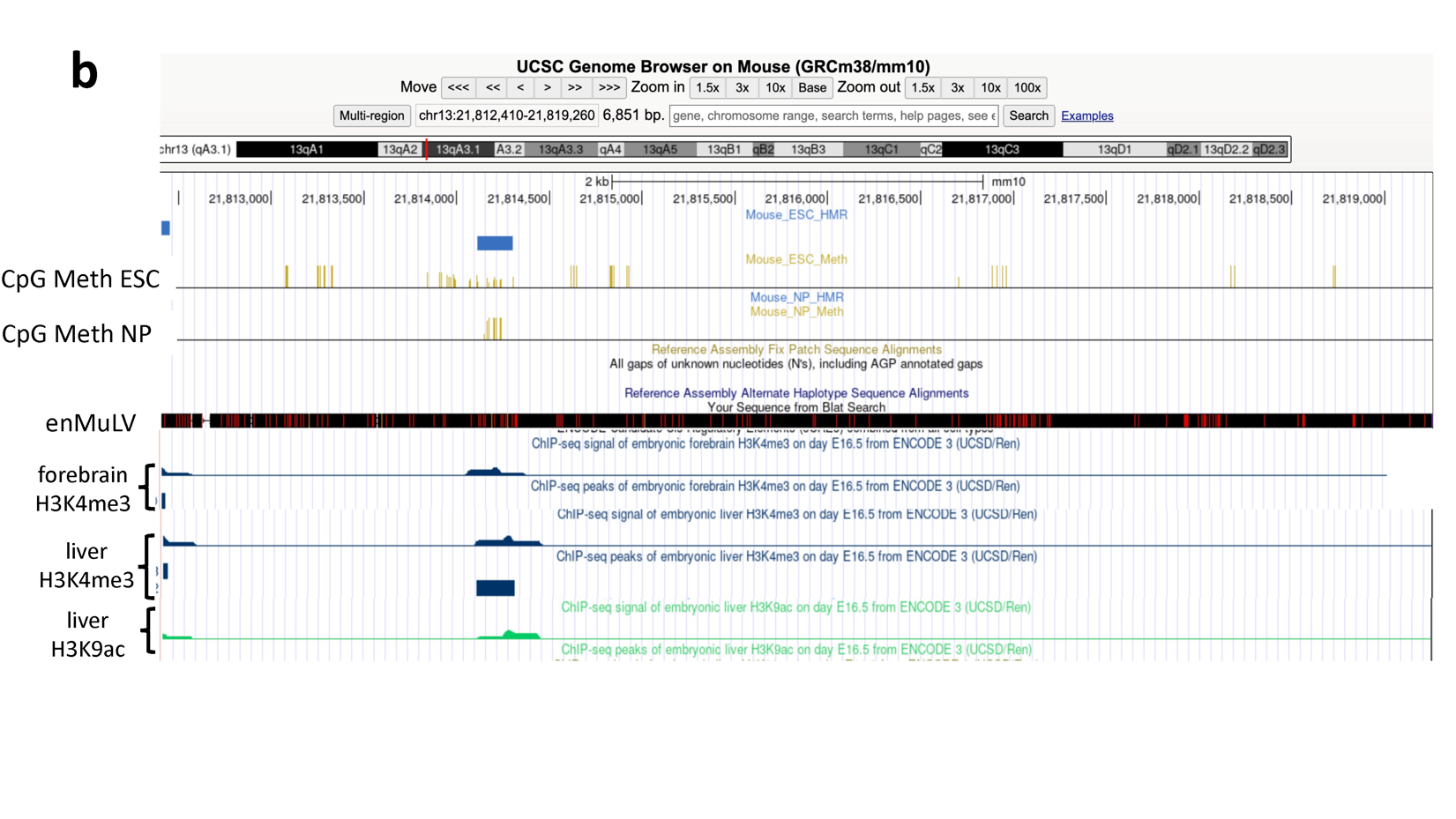

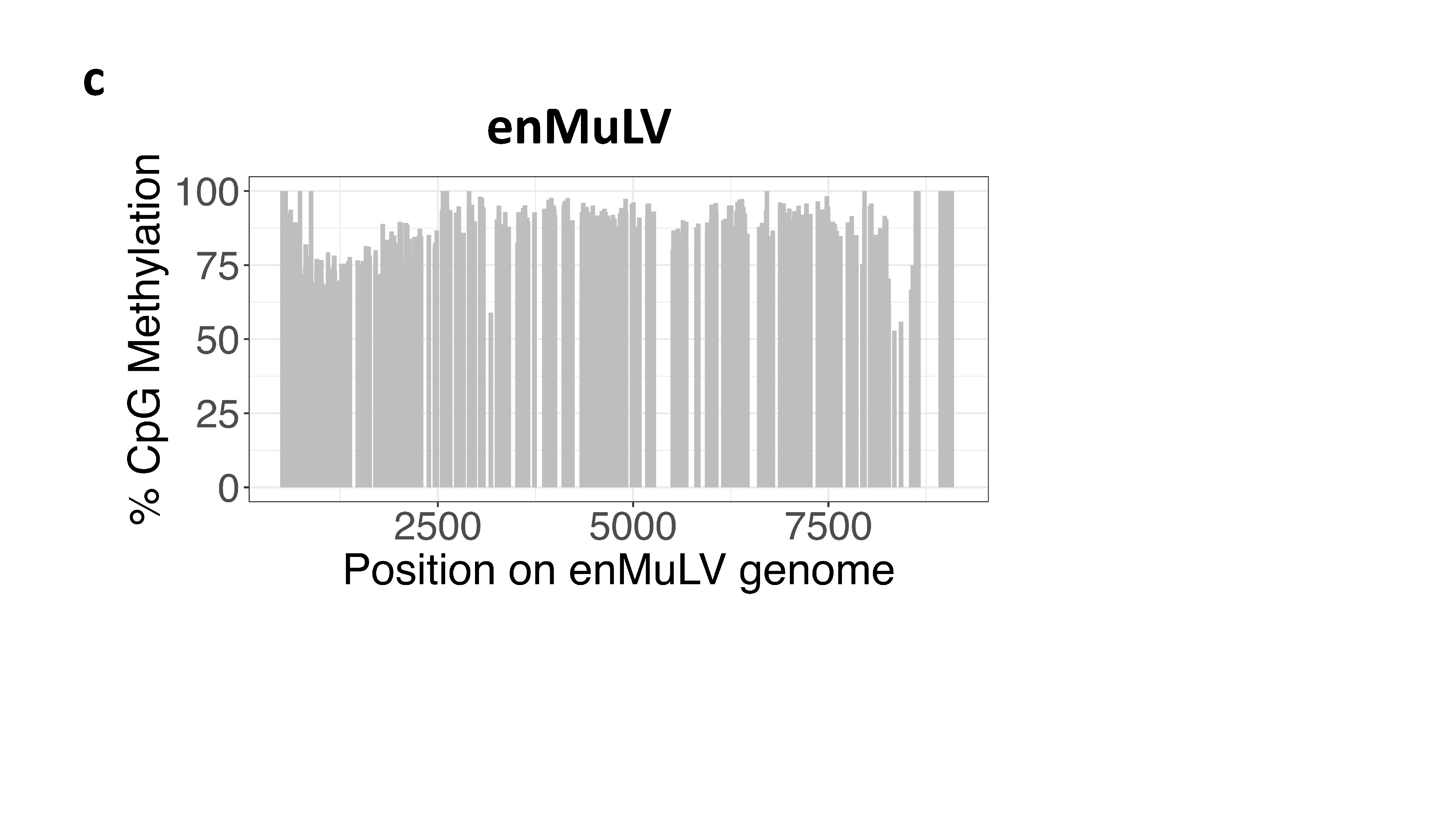

**Supplementary Figure S6:** CpG methylation in endogenous MuLV overlays histone modifications. **a)** enMuLV locus on Chromosome 5 exhibits H3K9me3 and H3K36me3 in the body of the provirus in Day 16 mouse embryo forebrain, with heavy CpG methylation across the entire provirus in neural progenitor cells. **b)** enMuLV locus on Chromosome 13 exhibits H3K4me3 and H3K9ac in Day 16 mouse embryonic forebrain and liver, with sparse CpG methylation across the provirus in both embryonic stem cells and neural progenitors. A hypomethylated peak is noted in embryonic stem cells, corresponding with a peak of CpG methylation in neural progenitors overlaying the site of these activation-associated histone modifications. **c)** Percent CpG methylation across the enMuLV provirus in bisulfite converted DNA from mouse liver tissue.

| **Supplementary Table S1:** Hypermut3 analysis of enMuLV. Shaded sequences meet threshold for significance. | | | | | | |
| --- | --- | --- | --- | --- | --- | --- |
| **seq_name** | **primary_matches** | **potential_primaries** | **control_matches** | **potential_controls** | **rate_ratio** | **fisher_p** |
| **Mpmv10** | 297 | 984 | 191 | 791 | 1.25 | 0.00267328 |
| **Mpmv11** | 6 | 882 | 1 | 1195 | 8.13 | 0.0259101 |
| **Mpmv12** | 362 | 1265 | 154 | 812 | 1.51 | 3.30E-07 |
| **Mpmv13** | 44 | 869 | 46 | 1191 | 1.31 | 0.113906 |
| **Mpmv2** | 362 | 1265 | 154 | 812 | 1.51 | 3.30E-07 |
| **Mpmv3** | 452 | 1495 | 123 | 582 | 1.43 | 1.48E-05 |
| **Mpmv4** | 0 | 883 | 2 | 1194 | 0 | 1 |
| **Mpmv5** | 229 | 794 | 274 | 1283 | 1.35 | 7.42E-05 |
| **Mpmv6** | 278 | 823 | 268 | 1254 | 1.58 | 2.99E-10 |
| **Mpmv7** | 10 | 882 | 0 | 1195 | inf | 0.00018514 |
| **Mpmv8** | 475 | 1537 | 126 | 540 | 1.32 | 0.00044308 |
| **Mpmv9** | 486 | 1592 | 135 | 485 | 1.1 | 0.140509 |
| **Pmv1** | 433 | 1434 | 155 | 640 | 1.25 | 0.00290729 |
| **Pmv10** | 233 | 792 | 220 | 941 | 1.26 | 0.00261461 |
| **Pmv11** | 340 | 1183 | 189 | 891 | 1.35 | 5.57E-05 |
| **Pmv12** | 436 | 1434 | 155 | 640 | 1.26 | 0.00216564 |
| **Pmv13** | 435 | 1435 | 155 | 639 | 1.25 | 0.00261842 |
| **Pmv14** | 343 | 1184 | 188 | 890 | 1.37 | 2.85E-05 |
| **Pmv15** | 435 | 1435 | 156 | 639 | 1.24 | 0.00330055 |
| **Pmv16** | 433 | 1434 | 157 | 640 | 1.23 | 0.0045706 |
| **Pmv17** | 217 | 733 | 218 | 930 | 1.26 | 0.00273339 |
| **Pmv18** | 429 | 1359 | 156 | 671 | 1.36 | 5.19E-05 |
| **Pmv19** | 428 | 1430 | 153 | 644 | 1.26 | 0.00207152 |
| **Pmv2** | 405 | 1288 | 199 | 785 | 1.24 | 0.00171338 |
| **Pmv20** | 281 | 926 | 255 | 1127 | 1.34 | 4.70E-05 |
| **Pmv21** | 437 | 1432 | 155 | 642 | 1.26 | 0.0016245 |
| **Pmv22** | 207 | 1065 | 96 | 1009 | 2.04 | 7.72E-11 |
| **Pmv23** | 345 | 1184 | 189 | 890 | 1.37 | 2.59E-05 |
| **Pmv24** | 432 | 1436 | 155 | 638 | 1.24 | 0.00381534 |
| **Pmv4** | 402 | 1273 | 176 | 801 | 1.44 | 1.04E-06 |
| **Pmv5** | 358 | 1245 | 183 | 829 | 1.3 | 0.00038546 |
| **Pmv6** | 431 | 1435 | 155 | 639 | 1.24 | 0.0038436 |
| **Pmv7** | 433 | 1435 | 155 | 639 | 1.24 | 0.0031778 |
| **Pmv8** | 351 | 1117 | 171 | 757 | 1.39 | 1.56E-05 |
| **Pmv9** | 434 | 1434 | 154 | 640 | 1.26 | 0.00208125 |
| **Xmv10** | 191 | 673 | 194 | 1009 | 1.48 | 8.87E-06 |
| **Xmv12** | 244 | 851 | 263 | 1198 | 1.31 | 0.00032594 |
| **Xmv13** | 246 | 814 | 258 | 1233 | 1.44 | 1.31E-06 |
| **Xmv15** | 238 | 782 | 315 | 1271 | 1.23 | 0.00306176 |
| **Xmv16** | 193 | 664 | 228 | 1040 | 1.33 | 0.00055747 |
| **Xmv17** | 196 | 641 | 222 | 986 | 1.36 | 0.00018512 |
| **Xmv18** | 224 | 810 | 271 | 1237 | 1.26 | 0.00183832 |
| **Xmv19** | 221 | 744 | 294 | 1202 | 1.21 | 0.00643881 |
| **Xmv41** | 251 | 836 | 281 | 1210 | 1.29 | 0.00035746 |
| **Xmv42** | 168 | 629 | 181 | 937 | 1.38 | 0.00038119 |
| **Xmv43** | 232 | 808 | 246 | 1238 | 1.44 | 2.84E-06 |
| **Xmv8** | 248 | 845 | 247 | 1206 | 1.43 | 2.74E-06 |
| **Xmv9** | 252 | 800 | 276 | 1247 | 1.42 | 1.69E-06 |

| **Supplementary Table S2:** SEA results for all MuLVs examined. | | | | | | | | |
| --- | --- | --- | --- | --- | --- | --- | --- | --- |
| **exMuLV** | | | | | | | | |
| **Motif** | **Location** | **Filter** | **P-value** | **E-value** | **Q-value** | **# in exMuLV** | **# in enMuLV** | **Enrichment Ratio** |
| H3K27ac, H3K9ac | CGI within Gag | 4214 | 1.54E-15 | 5.69E-13 | 3.09E-13 | 16/16 | 0/49 | 50 |
| H3K27ac | CGI within Gag | 2942 | 7.72E-14 | 2.85E-11 | 1.71E-12 | 15/16 | 0/49 | 47.1 |
| H3K9ac | CGI within Gag | 2979 | 7.72E-14 | 2.85E-11 | 1.71E-12 | 15/16 | 0/49 | 47.1 |
| **MPMV** | | | | | | | | |
| **Motif** | **Location** | **Filter** | **P-value** | **E-value** | **Q-value** | **# in MPMV** | **# in exMuLV** | **Enrichment Ratio** |
| H3K27ac | CGI in leader region | 963 | 3.80E-05 | 1.40E-02 | 1.70E-04 | 13/13 | 0/16 | 17.0 |
| H3K27ac | CGI in leader region | 3688 | 3.80E-05 | 1.40E-02 | 1.70E-04 | 13/13 | 0/16 | 17.0 |
| H3K27ac | CGI in leader region | 4252 | 3.80E-05 | 1.40E-02 | 1.70E-04 | 13/13 | 0/16 | 17.0 |
| H3K27me3 | CGI in leader region | 4252 | 3.80E-05 | 1.40E-02 | 1.70E-04 | 13/13 | 0/16 | 17.0 |
| H3K27me3 | CGI in leader region | 4343 | 3.80E-05 | 1.40E-02 | 1.70E-04 | 13/13 | 0/16 | 17.0 |
| H3K27me3 | CGI in leader region | 4321 | 3.80E-05 | 1.40E-02 | 1.70E-04 | 13/13 | 0/16 | 17.0 |
| H3K27me3 | Gag | 4399 | 3.80E-05 | 1.40E-02 | 1.70E-04 | 13/13 | 0/16 | 17.0 |
| H3K27me3 | LTR | 3682 | 3.80E-05 | 1.40E-02 | 1.70E-04 | 13/13 | 0/16 | 17.0 |
| H3K27me3 | Gag | 4194 | 3.80E-05 | 1.40E-02 | 1.70E-04 | 13/13 | 0/16 | 17.0 |
| H3K27me3 | CGI at pol/env junction | 2580 | 3.80E-05 | 1.40E-02 | 1.70E-04 | 13/13 | 0/16 | 17.0 |
| H3K27me3/H3K4me3 | CGI in leader region | 4403 | 3.80E-05 | 1.40E-02 | 1.70E-04 | 13/13 | 0/16 | 17.0 |
| H3K4me3/H3K9ac | CGI in leader region | 3898 | 3.80E-05 | 1.40E-02 | 1.70E-04 | 13/13 | 0/16 | 17.0 |
| H3K4me3 | CGI in leader region | 4043 | 3.80E-05 | 1.40E-02 | 1.70E-04 | 13/13 | 0/16 | 17.0 |
| **PMV** | | | | | | | | |
| **Motif** | **Location** | **Filter** | **P-value** | **E-value** | **Q-value** | **# in PMV** | **# in exMuLV** | **Enrichment Ratio** |
| H3K27me2/H3K27ac | CGI in leader region | 4252 | 6.28E-06 | 2.32E-03 | 3.43E-05 | 23/23 | 0/16 | 17.0 |
| H3K27me3 | CGI in leader region | 4343 | 6.28E-06 | 2.32E-03 | 3.43E-05 | 23/23 | 0/16 | 17.0 |
| H3K27me3/H3K9ac | CGI in leader region | 4410 | 6.28E-06 | 2.32E-03 | 3.43E-05 | 23/23 | 0/16 | 17.0 |
| H3K27me3 | CGI in leader region | 4405 | 6.28E-06 | 2.32E-03 | 3.43E-05 | 23/23 | 0/16 | 17.0 |
| H3K27me3 | CGI in leader region | 4419 | 6.28E-06 | 2.32E-03 | 3.43E-05 | 23/23 | 0/16 | 17.0 |
| H3K4me3 | CGI in leader region | 2506 | 6.28E-06 | 2.32E-03 | 3.43E-05 | 23/23 | 0/16 | 17.0 |
| H3K4me3/H3K9ac | CGI in leader region | 4128 | 6.28E-06 | 2.32E-03 | 3.43E-05 | 23/23 | 0/16 | 17.0 |
| H3K4me3 | CGI in leader region | 4223 | 6.28E-06 | 2.32E-03 | 3.43E-05 | 23/23 | 0/16 | 17.0 |
| H3K4me3 | CGI in leader region | 4348 | 6.28E-06 | 2.32E-03 | 3.43E-05 | 23/23 | 0/16 | 17.0 |
| H3K4me3 | CGI in leader region | 4480 | 6.28E-06 | 2.32E-03 | 3.43E-05 | 23/23 | 0/16 | 17.0 |
| H3K27ac | CGI in leader region | 963 | 6.28E-06 | 2.32E-03 | 3.43E-05 | 23/23 | 0/16 | 17.0 |
| H3K27ac | CGI in leader region | 3564 | 6.28E-06 | 2.32E-03 | 3.43E-05 | 23/23 | 0/16 | 17.0 |
| H3K27ac | CGI in leader region | 3688 | 6.28E-06 | 2.32E-03 | 3.43E-05 | 23/23 | 0/16 | 17.0 |
| H3K4me3 | CGI in leader region | 4043 | 6.28E-06 | 2.32E-03 | 3.43E-05 | 23/23 | 0/16 | 17.0 |
| H3K4me3/H3K9ac | CGI in leader region | 4053 | 6.28E-06 | 2.32E-03 | 3.43E-05 | 23/23 | 0/16 | 17.0 |
| H3K27ac/H3K9ac | CGI in leader region | 4606 | 6.28E-06 | 2.32E-03 | 3.43E-05 | 23/23 | 0/16 | 17.0 |
| H3K9ac | CGI in leader region | 4282 | 6.28E-06 | 2.32E-03 | 3.43E-05 | 23/23 | 0/16 | 17.0 |
| H3K9ac | CGI in leader region | 2478 | 6.28E-06 | 2.32E-03 | 3.43E-05 | 23/23 | 0/16 | 17.0 |
| H3K9me3 | Proximal provirus | 4620 | 6.28E-06 | 2.32E-03 | 3.43E-05 | 23/23 | 0/16 | 17.0 |
| H3K9me3 | Proximal provirus | 223 | 6.28E-06 | 2.32E-03 | 3.43E-05 | 23/23 | 0/16 | 17.0 |
| H3K9me3 | Proximal provirus | 2409 | 6.28E-06 | 2.32E-03 | 3.43E-05 | 23/23 | 0/16 | 17.0 |
| H3K9me3 | Proximal provirus | 2548 | 6.28E-06 | 2.32E-03 | 3.43E-05 | 23/23 | 0/16 | 17.0 |
| H3K9me3 | Proximal provirus | 3036 | 6.28E-06 | 2.32E-03 | 3.43E-05 | 23/23 | 0/16 | 17.0 |
| H3K9me3 | Proximal provirus | 706 | 6.28E-06 | 2.32E-03 | 3.43E-05 | 23/23 | 0/16 | 17.0 |
| H3K9me3 | Proximal provirus | 734 | 6.28E-06 | 2.32E-03 | 3.43E-05 | 23/23 | 0/16 | 17.0 |
| H3K27ac | 5' LTR | 1018 | 6.28E-06 | 2.32E-03 | 3.43E-05 | 23/23 | 0/16 | 17.0 |

| **Supplementary Table S3:** SEA results for all FeLVs examined. | | | | | | | | |
| --- | --- | --- | --- | --- | --- | --- | --- | --- |
| **enFeLV** | | | | | | | | |
| **Motif** | **Location** | **Cluster** | **P-value** | **E-value** | **Q-value** | **# in enFeLV** | **# in exFeLV** | **Enrichment Ratio** |
| None with E-value < 0.05 | | | | | | | | |
| **exFeLV** | | | | | | | | |
| **Motif** | **Location** | **Cluster** | **P-value** | **E-value** | **Q-value** | **# in exFeLV** | **# in enFeLV** | **Enrichment Ratio** |
| H3K4me3 | Proximal LTR | 24 | 7.77E-05 | 2.49E-02 | 5.43E-04 | 8/8 | 0/8 | 9.0 |
| H3K4me3 | CGI in pol/env junction | 103 | 7.77E-05 | 2.49E-02 | 5.43E-04 | 8/8 | 0/8 | 9.0 |
| H3K4me3 | Proximal LTR | 116 | 7.77E-05 | 2.49E-02 | 5.43E-04 | 8/8 | 0/8 | 9.0 |
| H3K4me3 | Proximal LTR | 125 | 7.77E-05 | 2.49E-02 | 5.43E-04 | 8/8 | 0/8 | 9.0 |
| H3K4me3 | CGI in proximal gag | 138 | 7.77E-05 | 2.49E-02 | 5.43E-04 | 8/8 | 0/8 | 9.0 |
| H3K4me3 | CGI in proximal gag | 185 | 7.77E-05 | 2.49E-02 | 5.43E-04 | 8/8 | 0/8 | 9.0 |
| H3K4me3 | Proximal LTR | 194 | 7.77E-05 | 2.49E-02 | 5.43E-04 | 8/8 | 0/8 | 9.0 |
| H3K27ac | CGI in U5 of LTR | 101 | 7.77E-05 | 2.49E-02 | 5.43E-04 | 8/8 | 0/8 | 9.0 |
| H3K27ac | CGI in proximal gag and 5' LTR | 110 | 7.77E-05 | 2.49E-02 | 5.43E-04 | 8/8 | 0/8 | 9.0 |
| H3K27ac | CGI in U5 of LTR and CGI in distal gag | 140 | 7.77E-05 | 2.49E-02 | 5.43E-04 | 8/8 | 0/8 | 9.0 |
| H3K27ac | Proximal LTR | 157 | 7.77E-05 | 2.49E-02 | 5.43E-04 | 8/8 | 0/8 | 9.0 |
| H3K27ac | Proximal LTR | 236 | 7.77E-05 | 2.49E-02 | 5.43E-04 | 8/8 | 0/8 | 9.0 |
| H3K27ac | CGI in proximal gag | 319 | 7.77E-05 | 2.49E-02 | 5.43E-04 | 8/8 | 0/8 | 9.0 |

| **Supplementary Table S4:** SEA results for all JSRVs examined. | | | | | | | | |
| --- | --- | --- | --- | --- | --- | --- | --- | --- |
| **enJSRV** | | | | | | | | |
| **Motif** | **Location** | **Cluster** | **P-value** | **E-value** | **Q-value** | **# in enJSRV** | **# in exJSRV** | **Enrichment Ratio** |
| H3K4me3 | CGI in U5 of LTR | 207 | 9.63E-12 | 3.08E-09 | 1.70E-09 | 26/26 | 1/20 | 10.5 |
| H3K4me3 | CGI in pol/env junction | 313 | 3.16E-10 | 1.01E-07 | 1.19E-08 | 23/26 | 0/20 | 18.7 |
| H3K4me3 | CGI in gag | 170 | 1.88E-09 | 6.02E-07 | 1.67E-08 | 24/26 | 1/20 | 9.7 |
| **exJSRV ENTV** | | | | | | | | |
| **Motif** | **Location** | **Cluster** | **P-value** | **E-value** | **Q-value** | **# in ENTV** | **# in enJSRV** | **Enrichment Ratio** |
| H3K4me3 | CGI in U5 of LTR | 10 | 6.50E-07 | 2.08E-04 | 8.88E-06 | 13/13 | 0/26 | 27.0 |
| H3K4me3 | CGI in pol/env junction | 109 | 6.50E-07 | 2.08E-04 | 8.88E-06 | 13/13 | 0/26 | 27.0 |
| H3K4me3 | CGI in pol/env junction | 281 | 6.50E-07 | 2.08E-04 | 8.88E-06 | 13/13 | 0/26 | 27.0 |
| H3K4me3 | CGI in pol/env junction | 94 | 3.25E-05 | 1.04E-02 | 1.94E-04 | 13/13 | 1/26 | 12.5 |
| H3K4me3 | Proximal LTR | 82 | 6.27E-06 | 2.01E-03 | 4.70E-05 | 13/13 | 1/26 | 13.5 |
| H3K27ac | CGI in pol/env junction | 2 | 6.50E-07 | 2.08E+04 | 1.04E-05 | 13/13 | 0/26 | 27.0 |
| H3K27ac | CGI in U5 of LTR | 69 | 6.50E-07 | 2.08E+04 | 1.04E-05 | 13/13 | 0/26 | 13.3 |
| H3K27ac | Proximal LTR | 163 | 1.94E-06 | 6.22E-04 | 2.73E-05 | 12/13 | 0/26 | 25.1 |
| H3K27ac | CGI in U5 of LTR | 191 | 3.25E-05 | 1.04E-02 | 2.28E-04 | 13/13 | 1/26 | 9.0 |
| **exJSRV OPP** | | | | | | | | |
| **Motif** | **Location** | **Cluster** | **P-value** | **E-value** | **Q-value** | **# in OPP** | **# in enJSRV** | **Enrichment Ratio** |
| H3K4me3 | CGI in pol/env junction | 29 | 9.40E-05 | 3.01E-02 | 2.16E-03 | 6/7 | 0/26 | 11.1 |
| H3K4me3 | U5 of LTR | 311 | 9.40E-05 | 3.01E-02 | 2.16E-03 | 6/7 | 0/26 | 11.1 |
| H3K4me3 | CGI in gag | 276 | 2.00E-05 | 6.41E-03 | 2.16E-03 | 7/7 | 0/26 | 13.5 |
| H3K27ac | Proximal LTR | 83 | 2.00E-05 | 6.41E-03 | 2.50E-03 | 7/7 | 0/26 | 27.0 |
| H3K27ac | CGI in gag | 242 | 9.40E-05 | 3.01E-02 | 2.50E-03 | 6/7 | 0/26 | 23.6 |
